# Genome compartments guide protamine replacement and genome stability during spermiogenesis

**DOI:** 10.64898/2026.03.08.710361

**Authors:** Masashi Hada, Chen Zhong, Erina Inoue, Yuko Fukuda, Chizuko Koga, Mihoko Hosokawa, Shin-ichiro Chuma, Shoko Sato, Hitoshi Kurumizaka, Masahito Ikawa, Sung Hee Baek, Yuki Okada

## Abstract

Compartment-scale genome organisation persists in mammalian sperm, yet how histone–protamine replacement is orchestrated in space and time during spermiogenesis remains unclear. Here we combine stage-resolved purification of mouse spermatids with spike-in-normalised ATAC-seq and PRM1 CUT&Tag to map chromatin accessibility and protamine incorporation across spermiogenesis. We uncover a transient, genome-wide hyper-accessible phase coincident with replacement that is uncoupled from transcription and suppressed in catalytic PHF7-mutant spermatids. PRM1 loading initiates within accessible A-compartment chromatin and later spreads across both A- and B-compartments, whereas protamine-null mice demonstrated that PRM1/ PRM2 deficiency selectively destabilises A-compartment closure. Sequencing of short DNA fragments from Prm1/Prm2 dosage-reduced epididymal sperm reveals early enrichment of breakage at protamine-targeted A-compartment regions, which dissipates as fragmentation becomes genome-wide during epididymal transit. Together, our data place histone–protamine replacement within a compartment-centred framework that links 3D genome domains to the timing, targeting and integrity of the sperm genome.

## Introduction

Packaging of the paternal genome into an exceptionally compact nucleus is the defining chromatin transformation of spermiogenesis^1,2^. During this transition, canonical nucleosomes are evicted and replaced by transition nuclear proteins (TNPs) and, ultimately, protamines (PRMs), yielding a highly condensed chromatin that safeguards the genome and supports fertility^3,4^. Thus, perturbations of this replacement result in male infertility and sperm DNA damage, underscoring that histone eviction and PRM incorporation are essential, tightly regulated steps rather than a passive end-stage of differentiation^4^.

At the molecular level, histone-to-protamine replacement is preceded by a wave of histone hyperacetylation in elongating spermatids, which is thought to destabilise nucleosomes and prime chromatin for large-scale remodelling^5^. A key reader of this acetylation is the testis-specific BET protein BRDT, which binds acetylated H4 and promotes acetylation-dependent chromatin reorganisation and histone eviction^5^. Ubiquitin signaling provides an additional layer of control: PHF7 has been shown to ubiquitylate histones (including H2A and H3) and to facilitate BRDT-dependent eviction^6,7^. Downstream of eviction, hyperacetylated histones can be eliminated via acetylation-dependent proteasomal degradation mediated by the PA200/PSME4 proteasome activator^8^. In parallel, testis-specific histone variants and intermediates—including H2A.L.2, together with TNP1/TNP2—create transient chromatin states that facilitate ordered protamine assembly and terminal compaction^9^.

However, despite these advances, the temporal order and spatial logic of this replacement remain incompletely defined. While classical histological staging provides a robust morphological framework, translating spermiogenesis into genome-wide, step-resolved maps has been challenging. Recent flow cytometry-based cell sorting protocols have enabled highly enriched isolation of germ-cell populations and coarse subdivision of spermatids, yet they do not readily visualise spermiogenesis as a continuous maturation trajectory from which arbitrary steps can be isolated for downstream genomic assays^10^. It therefore remains unclear whether condensation and protamine deposition proceed uniformly across the genome or are biased towards specific chromatin domains, how histone eviction is coupled to subsequent protamination in vivo, and how any such spatial bias relates to higher-order genome organisation.

Hi-C analyses provide a complementary view of sperm genome organisation. Bulk Hi-C studies have detected A/B compartmentalisation and, in some datasets, TAD- and loop-like features in mouse sperm^11,12^, but related work in mouse and human sperm likewise reported prominent compartment signals despite limited evidence for obvious TADs^13–15^. However, reports of TAD- and loop-like features have been inconsistent, in part owing to differences in purification strategies and potential confounding by somatic or cell-free chromatin^16^. Recent advanced single-sperm reconstructions reconcile these discrepancies by showing that chromosome territories and A/B compartments are reproducibly detectable in both mouse and human sperm, whereas canonical TADs and loops are not evident at the population level^17^. This compartment-scale organisation therefore provides a natural framework for asking whether histone eviction and protamine loading proceed uniformly across the genome or are biased towards specific chromatin domains.

Here we combine spermiogenic step-resolved isolation of mouse spermatids with spike-in-normalised ATAC-seq and CUT&Tag to map chromatin accessibility and protamine incorporation across late spermiogenesis. This approach allows us to delineate the spatio-temporal sequence that couples histone eviction to protamine deposition and to place these events within a compartmental framework of the genome. This enables us to define a conceptual spermiogenic programme in which transient chromatin loosening precedes and enables region-selective protamine-driven compaction.

## Results

### Spermatids undergo genome-wide chromatin opening during histone-protamine replacement

We employed flow cytometry–based spermatid purification from histone H4–Venus/H3.3–mCherry double-transgenic mouse testes^18^. Seven fractions (P1–P7) of spermatids were gated and isolated (Fig. 1a, b). This strategy enables stepwise purification of spermatids based on the expression timing and abundance of H4–Venus and H3.3–mCherry. Notably, P4, where fluorescence shifts from H4–Venus dominance to H3.3–mCherry dominance, coincides with the histone-to-protamine (H–P) replacement window^18^. The corresponding spermiogenic steps represented by each gate, together with nuclear morphology and reporter expression, are shown in Fig. 1b, c.

**Figure 1.**
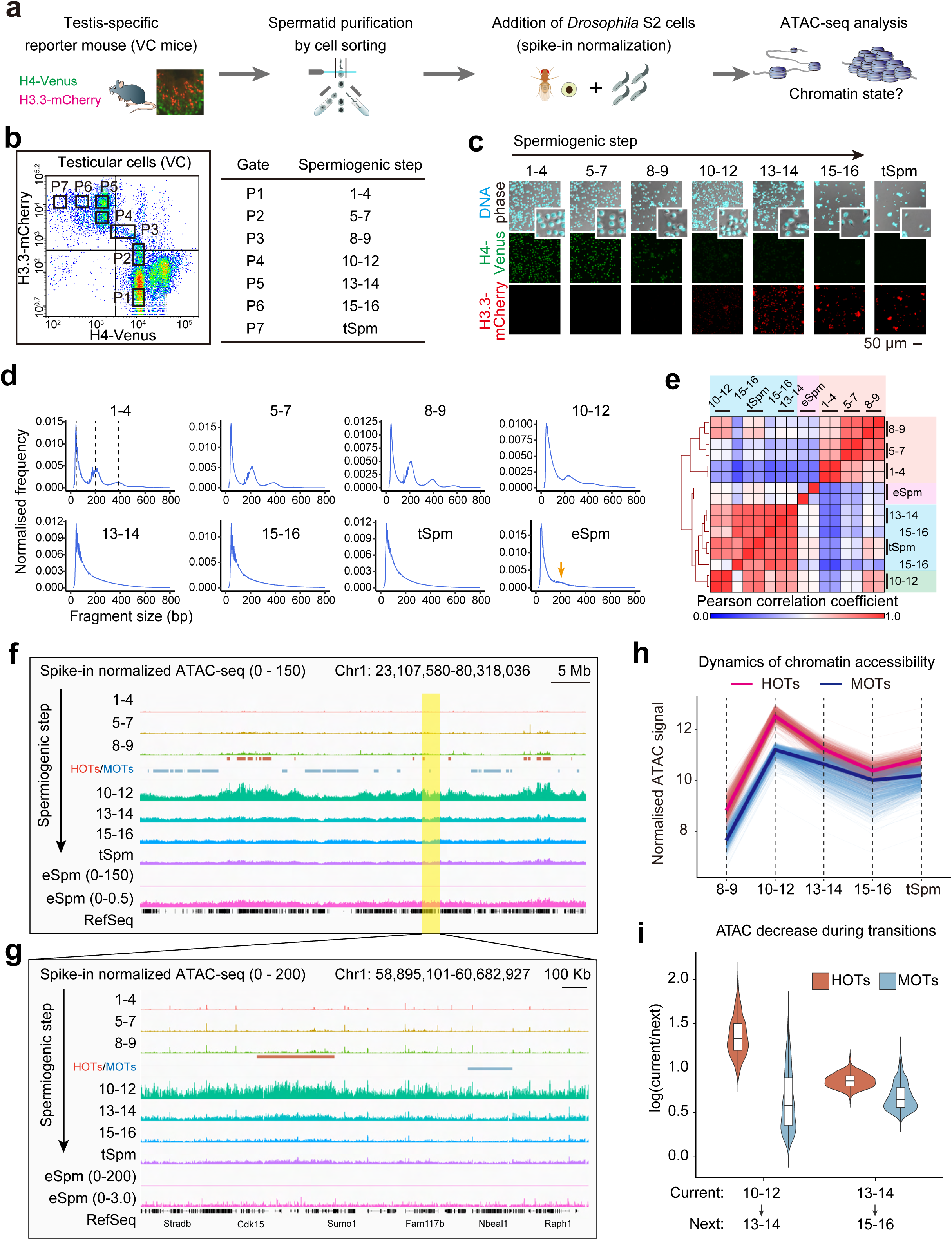
Global chromatin accessibility is extensively remodeled during spermiogenesis. **a**, Schematic overview of the spike-in–normalized ATAC-seq workflow. Spermatids were purified by cell sorting from testis-specific reporter mice expressing H4-Venus and H3.3-mCherry (VC). Drosophila S2 cells were added as an internal spike-in control prior to ATAC-seq library preparation. **b,** Cell sorting plot showing purification of spermatid populations from testicular cell suspensions. Gates corresponding to distinct spermiogenic steps are indicated. **c,** Morphological validation of sorted spermatids. Representative fluorescence microscopy images showing DNA, H4-Venus, and H3.3-mCherry signals across spermiogenic steps are shown. **d,** Fragment size distributions of ATAC-seq libraries across spermiogenic stages. The positions of nucleosome-free fragments and mono- and di-nucleosomal peaks are indicated by dashed lines in the step 1–4 spermatid panel. The nucleosomal ladder pattern progressively diminishes during mid-to-late spermiogenesis and reappears in epididymal sperm (yellow arrow). **e,** Heatmap showing Pearson correlation coefficients of genome-wide ATAC-seq signals across spermiogenic steps and epididymal sperm. Rows and columns are ordered by hierarchical clustering. **f, g,** Genome browser snapshots showing spike-in–normalized ATAC-seq signals across spermiogenic stages. Genomic regions corresponding to highly open territories (HOTs) and moderately open territories (MOTs) are indicated. The region shown in (**g)** represents a magnified view of the highlighted region in (**f)**. **h,** Stage-resolved dynamics of spike-in–normalized ATAC-seq signals within HOTs and MOTs. Individual lines represent one territory, and thick lines indicate median values. **i,** Quantification of stage-to-stage accessibility changes. The ratio of spike-in–normalized ATAC-seq signals between consecutive stages (“current” versus “next”) is shown for HOTs and MOTs. Abbreviations: tSpm, testicular sperm; eSpm, epididymal sperm.

Sorted cells were subjected to spike-in-normalised ATAC-seq. Fragment-length distributions of Tn5-digested DNA showed a typical nucleosome ladder up to steps 10-12, whereas the ladder became indistinct at steps 13-14. At later steps, the ladder disappeared, and reads were dominated by subnucleosomal fragments (<100 bp) (Fig. 1d, S1a). In epididymal sperm (eSpm), a minor mono-nucleosomal peak was detectable (Fig. 1d, arrow). Correlation analysis captured gradual changes in chromatin accessibility during spermiogenesis and separated samples collected up to steps 10-12 from those collected from steps 13-14 onwards (Fig. 1e). At representative loci, chromatin accessibility increased genome-wide at steps 10-12, coinciding with the timing of H–P replacement. However, this increase was not uniform: accessibility varied across regions spanning hundreds of kilobases to several megabases (Fig. 1f). We therefore stratified the genome into Hyper-Open Territories (HOTs), Intermediate-Open Territories (IOTs) and Moderately-Open Territories (MOTs) based on accessibility levels at steps 10–12 (Fig. 1f). HOTs and MOTs tended to correspond to higher and lower gene densities, respectively, although accessibility in MOTs was still elevated relative to steps 8–9 (Fig. 1f). Consistent with these domain-level properties, repeat-class analysis revealed enrichment of SINE elements in HOTs and of LINE elements in MOTs (Fig. S1b, S1c). At higher resolution, promoter-proximal peaks characteristic of somatic-like chromatin were evident at steps 1–4, 5–7 and 8–9 (Fig. 1g, S1d). Notably, HOTs showed a rise in baseline accessibility beginning at steps 8–9, whereas MOTs remained comparatively closed (Fig. 1g). Comparison of HOT and MOT accessibility across stages further demonstrated that HOT showed transient sharp chromatin openings from steps 8-9 to 10-12, followed by marked compaction towards steps 13-14, approaching MOT levels (Fig. 1h, 1i, S1e). In contrast, MOT exhibited relaxation from steps 8-9 to 10-12 with lower extent than HOT, with subsequent compaction being moderate (Fig. 1h, 1i, S1e). From steps 13–14 onwards, chromatin accessibility progressively declined genome-wide and became minimal in eSpm (Fig. 1f, 1g). Collectively, these ATAC-seq data reveal a transition from a somatic-like accessibility landscape to highly compacted sperm chromatin via a transient, genome-wide opening during H–P replacement.

### Chromatin accessibility correlates with the A/B compartmentalisation

Striking correlations between chromatin accessibility and gene density during spermiogenesis prompted us to examine the relationship between our ATAC-seq data and higher-order genome organisation. While the presence of TADs in condensed spermatids and sperm remains debated, A/B compartments have been consistently detected in sperm Hi-C studies^13–17^. Interestingly, A/B compartment identity in round spermatids correlated with the HOT/MOT pattern in both genomic distribution and signal intensity^19^ (Fig. 2a–c). Compartment-stratified differences in accessibility remained evident through steps 10–12 but became progressively less distinguishable thereafter as genome-wide compaction reduced ATAC-seq signal across both compartments (Fig. 2a, d).

**Figure 2.**
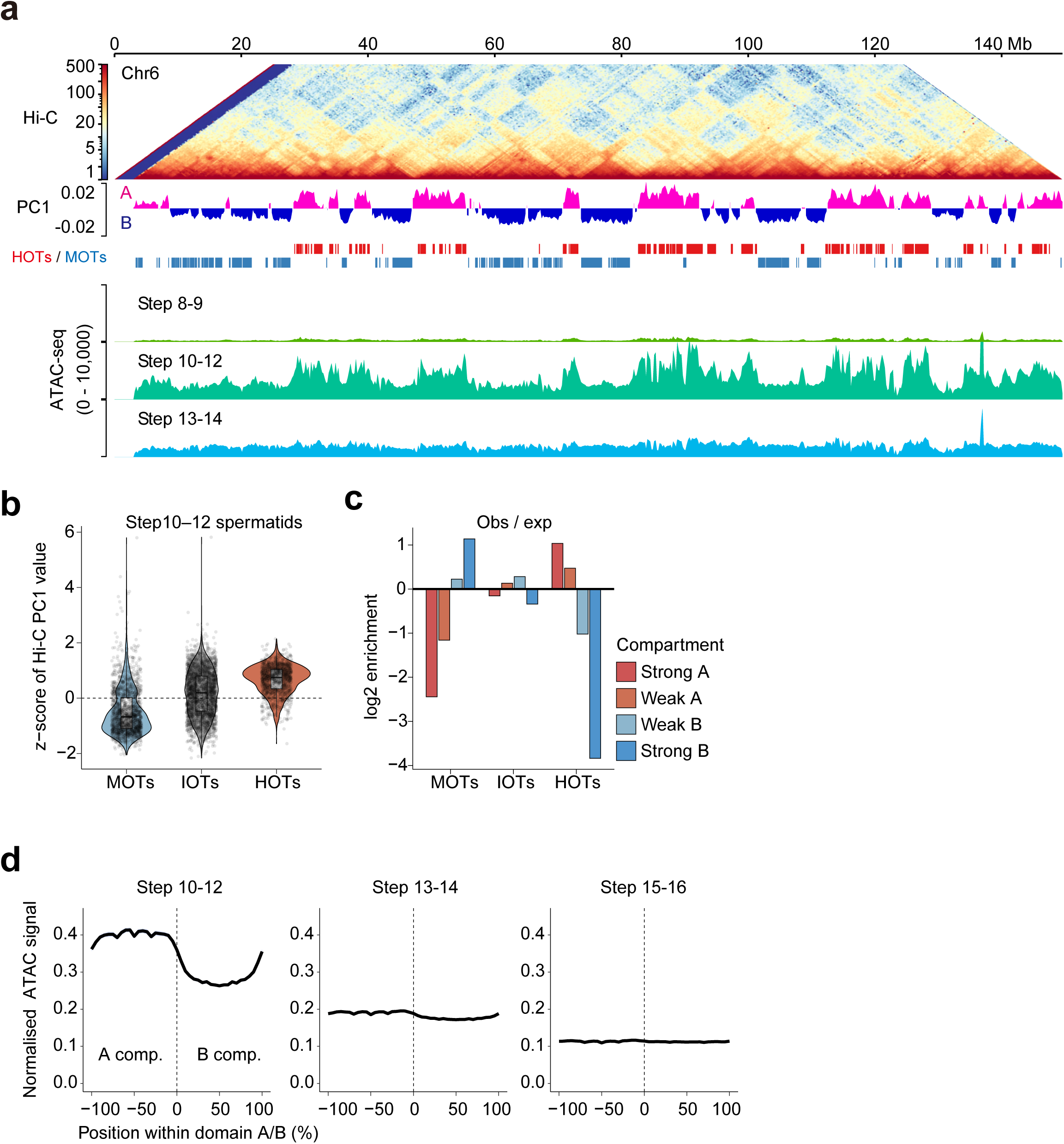
Chromatin accessibility territories are aligned with A/B compartment architecture during spermiogenesis. **a**, Integrated view of chromatin accessibility and three-dimensional genome organization across spermiogenic stages. A Hi-C contact map of mouse chromosome 6 in round spermatids^19^ is shown together with the first principal component (PC1) of the Hi-C correlation matrix, representing A/B compartment status. Hi-C data were obtained from GSE119805^19^. Genomic regions classified as HOTs and MOTs are indicated. Spike-in–normalized ATAC-seq signal tracks for step 8–9, step 10–12, and step 13–14 spermatids are shown below. **b,** Distribution of z-scored Hi-C PC1 values for HOTs, IOTs, and MOTs in step 10–12 spermatids. **c,** Enrichment of HOTs, IOTs, and MOTs within Hi-C compartments, shown as observed/expected log₂ enrichment. Compartments were stratified into strong A, weak A, weak B, and strong B based on PC1 values. **d,** Average spike-in–normalized ATAC-seq signal profiles across A/B compartment boundaries at different spermiogenic stages. Spike-in normalised ATAC-seq signal is plotted as a function of genomic distance from A/B compartment boundaries, with negative and positive values corresponding to positions within A and B compartments, respectively.

### Transient chromatin opening state is independent from transcription

We next performed RNA-seq to test whether these changes in chromatin accessibility correlate with transcription in spermatids. The same set of sorted populations used for ATAC-seq (except eSpm) was subjected to RNA-seq. PCA revealed a continuous trajectory of transcriptional state changes from steps 1–4 to testicular sperm (tSpm) (Fig. S2a). We detected 5,926 transcribed genes and classified them into three clusters based on expression dynamics (Fig. S2b). Cluster 1 (n = 3,973) peaked at steps 1–4 and gradually declined, with a re-increase in tSpm. Cluster 2 (n = 1,019) declined after step 9. Cluster 3 (n = 934) showed low expression at steps 1–4 and, once increased, remained comparatively stable thereafter. In gene ontology (GO) analysis, sperm-motility-related terms were significantly enriched: Cluster 1 was enriched for mitochondrial membrane, Cluster 2 for CatSper and cilium-related terms, and Cluster 3 for flagellum/cilium components (Fig. S2c). Notably, Cluster 2 was also enriched for the Cullin–RING ubiquitin ligase complex, consistent with transcription of components that may support extensive proteolysis following H–P replacement^20^.

We then compared ATAC-seq signal around transcription start sites (TSSs) for each cluster across spermiogenic steps. A typical positive relationship between cluster expression and TSS accessibility was evident up to steps 8-9. However, from steps 10-12 onwards, this relationship was lost (Fig. S2d). These results suggest that the transient genome-wide chromatin opening coincident with H–P replacement is uncoupled from transcriptional activation.

### Transient chromatin opening is missing in Phf7 catalytic mutant with altered H–P replacement

Because defective histone–protamine (H–P) replacement can cause excessive histone retention and male infertility^4^, we asked whether the accessibility trajectory is altered in a model with impaired histone eviction. We therefore examined a catalytic mutant of PHF7 (*Phf7^CA/CA^*), an E3 ligase targeting histones H2A and H3^6,7^. Previous studies showed that PHF7-dependent histone ubiquitylation facilitates histone eviction by modulating BRDT stability, and that *Phf7^CA/CA^* males are infertile owing to aberrant histone retention and subsequent sperm DNA fragmentation^7^. When H4–Venus/H3.3–mCherry transgenes were introduced into *Phf7^CA/CA^* mice, we observed reduced H3.3–mCherry incorporation after H–P replacement without apparent cell population corresponding to steps 13-14 (Fig. 3a, S3a). Instead, the cell population at the position corresponding to steps 15-16 in the wild type increased in *Phf7^CA/CA^* mice (Fig. 1b, 3a). Therefore, in subsequent experiments, this cell population was compared with steps 13-14 cells in *Phf7^CA/+^* control mice.

**Figure 3.**
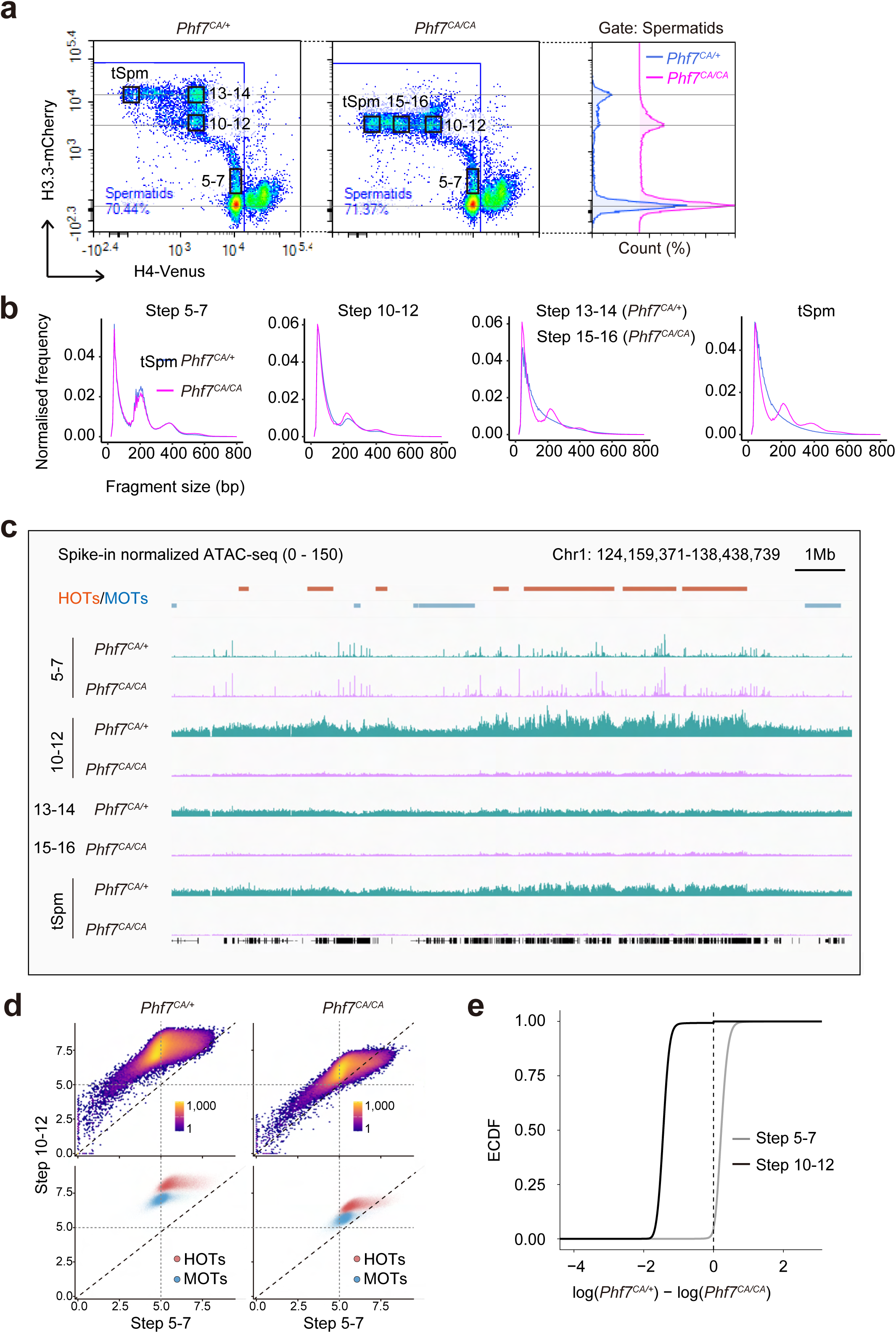
Impaired histone eviction suppresses hyper-opening of chromatin at step 10–12 of spermiogenesis. **a**, Cell sorting plots and gate settings showing spermatid populations isolated from control (*Phf7^CA/+^*) and catalytic mutant (*Phf7^CA/CA^*) mice. H4-Venus and H3.3-mCherry signals were used to gate spermatids, and representative fluorescence intensity distributions within the spermatid gate are shown on the right. **b,** Fragment size distributions of ATAC-seq libraries from *Phf7^CA/+^* (blue) and *Phf7^CA/CA^* (magenta) spermatids at the indicated spermiogenic stages. **c,** Genome browser snapshots showing spike-in–normalized ATAC-seq signals across spermiogenic stages in *Phf7^CA/+^* and *Phf7^CA/CA^* spermatids. Genomic regions corresponding to HOTs and MOTs are indicated. **d,** Scatter plots comparing log₂-transformed ATAC-seq signals between *Phf7^CA/+^* and *Phf7^CA/CA^*spermatids at steps 5–7 and steps 10–12. Regions corresponding to HOTs and MOTs are highlighted in the lower panels. **e,** Empirical cumulative distribution functions (ECDFs) showing differences in chromatin accessibility between *Phf7^CA/+^* and *Phf7^CA/CA^* spermatids at the indicated stages, calculated as log₂(*Phf7^CA/+^*) − log₂(*Phf7^CA/CA^*).

Fragment-length distributions revealed retained nucleosomal ladders in steps 15–16 spermatids and tSpm in the *Phf7^CA/CA^* mice (Fig. 3b). These observations were consistent with previous histopathology and immunostaining demonstrating excess histone retention in mutant spermatids^7^. Importantly, after showing the somatic-like, TSS-centered peak distribution at steps 5–7, *Phf7^CA/CA^* spermatids failed to exhibit genome-wide opening at steps 10–12 (Fig. 3c). As reported previously, we also confirmed comparable PRM1 expression in *Phf7^CA/CA^* and control testes by immunostaining^7^ (Fig. S3b). Correlation analysis demonstrated that both HOTs and MOTs in steps 10–12 were less accessible in *Phf7^CA/CA^*, whereas such differences were not observed in steps 5–7, in which both *Phf7^CA/+^* and *Phf7^CA/CA^* exhibited somatic-like peak patterns (Fig. 3c–e). Compartment-stratified differences in accessibility indicated that, in steps 10-12, *Phf7^CA/+^* display elevated accessibility within A-compartments, whereas the accessibility is globally reduced in *Phf7^CA/CA^* with less distinguishable compartmentalization (Fig. S3c, left panels). In steps 13-14, *Phf7^CA/+^* exhibited indistinct pattern similar to the wild type (Fig. 2d), while the compartment pattern is largely unchanged in *Phf7^CA/CA^* (Fig. S3c, right panels).

### PRM1 preferentially targets high-accessible chromatin in the A-compartment

These results from *Phf7^CA/CA^* mutants suggest that transient genome-wide chromatin opening is necessary for H–P replacement, and that productive incorporation of PRMs into a largely histone-depleted regions is required for robust chromatin compaction that prevents DNA fragmentation. We therefore profiled PRM1 incorporation across spermiogenesis by CUT&Tag to determine how protamine loading relates to chromatin accessibility.

Consistent with previous immunodetection, PRM1 signal first became prominent at steps 10–12 (Fig. 4a)^7^ and appeared in CUT&Tag profiles as dispersed kilobase-scale enrichment across the genome, with only modest compartmental preferences at this stage (Fig. 4b). By steps 13-14, PRM1 signal became organized into discrete megabase-scale regions that preferentially aligned with the A-compartment defined by Hi-C track; we termed these regions “<u>P</u>RM1 <u>p</u>referentially <u>a</u>ssociated <u>p</u>artitions (PPAPs) “ (Fig. 4b, c). PPAPs, HOTs and MOTs were detected across all autosomes without apparent preference to specific chromosomes (Fig. S4a). At 50-kb resolution, PPAPs overlapped extensively with HOTs but only minimally with MOTs (Fig. 4c), and were significantly enriched in HOTs and A-compartment bins particularly in strongly A-compartment region, while being depleted from MOTs and B-compartment bins (Fig. 4d). Importantly, PRM1 enrichment in steps 13–14 showed a graded relationship with the magnitude of accessibility loss: loci exhibiting the strongest decrease in ATAC-seq signal from steps 10–12 to 13–14 displayed the highest PRM1 enrichment, whereas loci with little or no decrease showed minimal enrichment (Fig. 4e). Together, these results support a model in which PRM1 is first detectable during the replacement window as dispersed binding, then becomes compartment-structured into A-enriched PPAPs as chromatin closes. In tSpm, PRM1 occupancy extended beyond PPAP boundaries and became more widespread across the genome (Fig. 4b). Concordantly, PRM1-high regions in tSpm were relatively enriched within B-compartment/MOT regions (Fig. S4b), and the graded coupling to accessibility change observed at steps 13–14 was not as evident at the tSpm stage (Fig. S4c).

**Figure 4.**
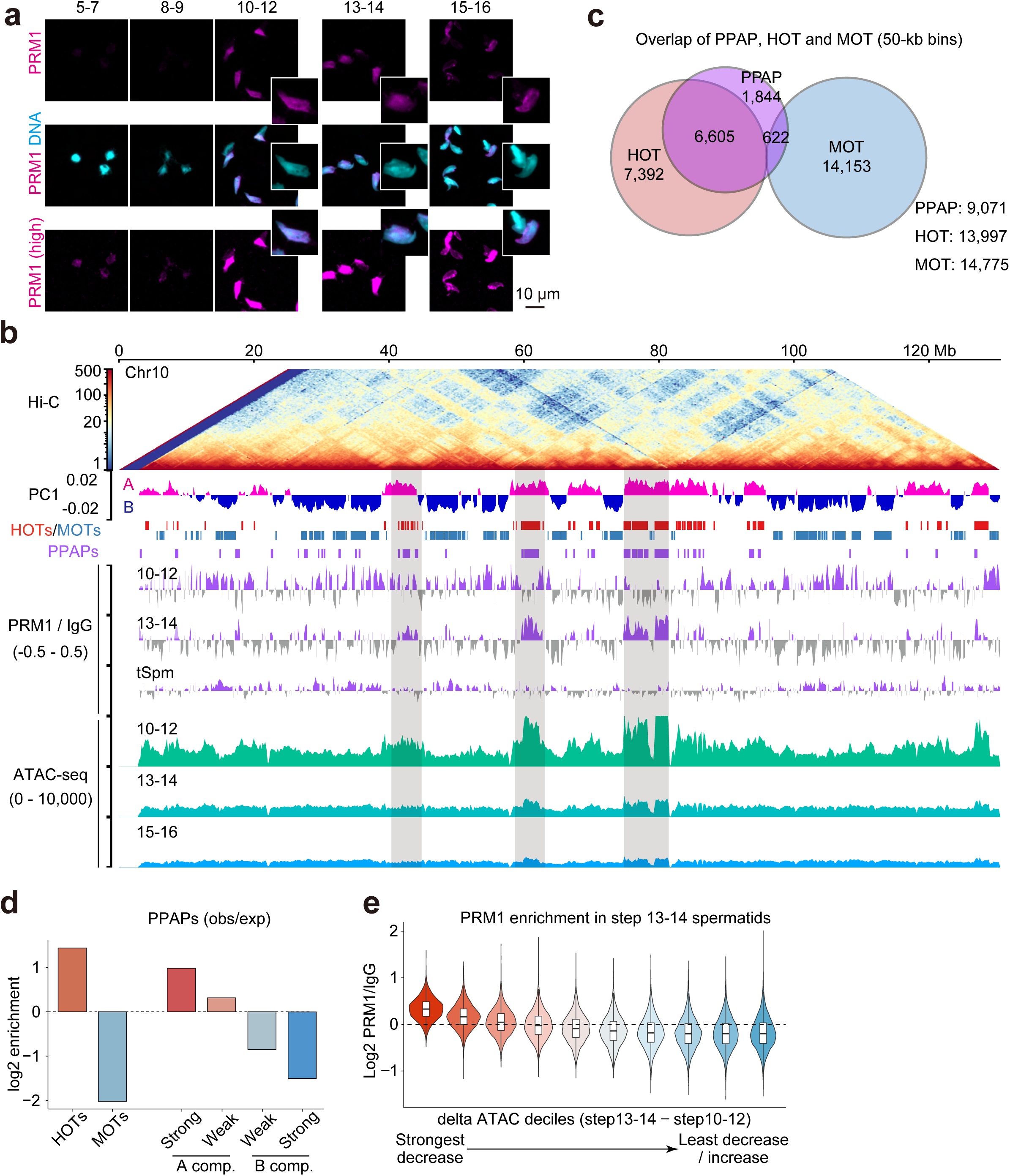
PRM1 incorporation is biased toward hyper-accessible chromatin territories during spermiogenesis. **a**, Immunofluorescence analysis of PRM1 (magenta), and DNA (cyan) in representative spermatogenic cells. Scale bar, 10 μm. **b**, Integrated genome-wide view of three-dimensional genome organization, chromatin accessibility, and PRM1 incorporation during spermiogenesis. A Hi-C contact map of mouse chromosome 10 is shown together with the PC1 value of the Hi-C correlation matrix, indicating A/B compartment status. HOTs and MOTs and PRM1 preferentially associated partitions (PPAPs) are indicated. PRM1 CUT&Tag signal (normalized to IgG) and spike-in–normalized ATAC-seq signal tracks for steps 10–12, steps 13–14, and tSpm are shown. **c**, Overlap between PPAPs, HOTs, and MOTs defined in 50-kb genomic bins. PPAPs were defined as genomic bins exhibiting significant PRM1 CUT&Tag enrichment in steps 13–14 spermatids. Numbers indicate bin counts for each category and their intersections. **d,** Observed/expected (obs/exp) enrichment of PPAPs within HOTs, MOTs, and A/B compartment subclasses (strong A, weak A, strong B, weak B) in step 13–14 spermatids. **e**, PRM1 enrichment stratified by the magnitude of chromatin accessibility decrease. Genomic bins were ranked according to the change in spike-in–normalized ATAC-seq signal between steps 10–12 and steps 13–14 (deltaATAC = steps13–14 − steps10–12) and grouped into deciles. Violin plots show PRM1 CUT&Tag enrichment (log₂ PRM1/IgG) in steps 13–14 spermatids for each decile.

### PRMs maintain the compacted chromatin state in the A-compartment

To further confirm the preference of protamines to A-compartment regions, we generated *Prm1/Prm2* double-knockout (dKO) mice and tested whether protamines are required to establish or maintain A-compartment compaction. *Prm1* and *Prm2* reside on mouse chromosome 16, ∼5 kb apart (Fig. 5a). Two sgRNAs targeting exon 1 of *Prm1* and *Prm2* were used to delete both genes simultaneously by CRISPR–Cas9, resulted in *Prm1/Prm2* mutant allele (Fig. 5a). Because *Prm1/Prm2*-mutant males are infertile in the heterozygous state, *Prm1/Prm2* dKO mice could not be generated by conventional mating, thus we performed <u>el</u>ongated <u>s</u>permatid injection (ELSI) using *Prm1/Prm2* double-heterozygous knock-out (dHET) spermatids and oocytes, respectively. To our knowledge, this is the first *Prm1/Prm2* dKO mice that have been ever reported, and all downstream analyses were performed using the homozygous dKO mice (i.e., not chimeras/mosaic founders), enabling the isolation of PRM1/PRM2-null spermatids. The deletion was confirmed by genomic PCR (Fig. S5a) and by loss of PRM1 and PRM2 immunoreactivity in the sorted spermatids and testis sections, respectively (Fig. 5b, S5b). Increased TNP1 was also observed in the dKO testis sections (Fig. S5c).

**Figure 5.**
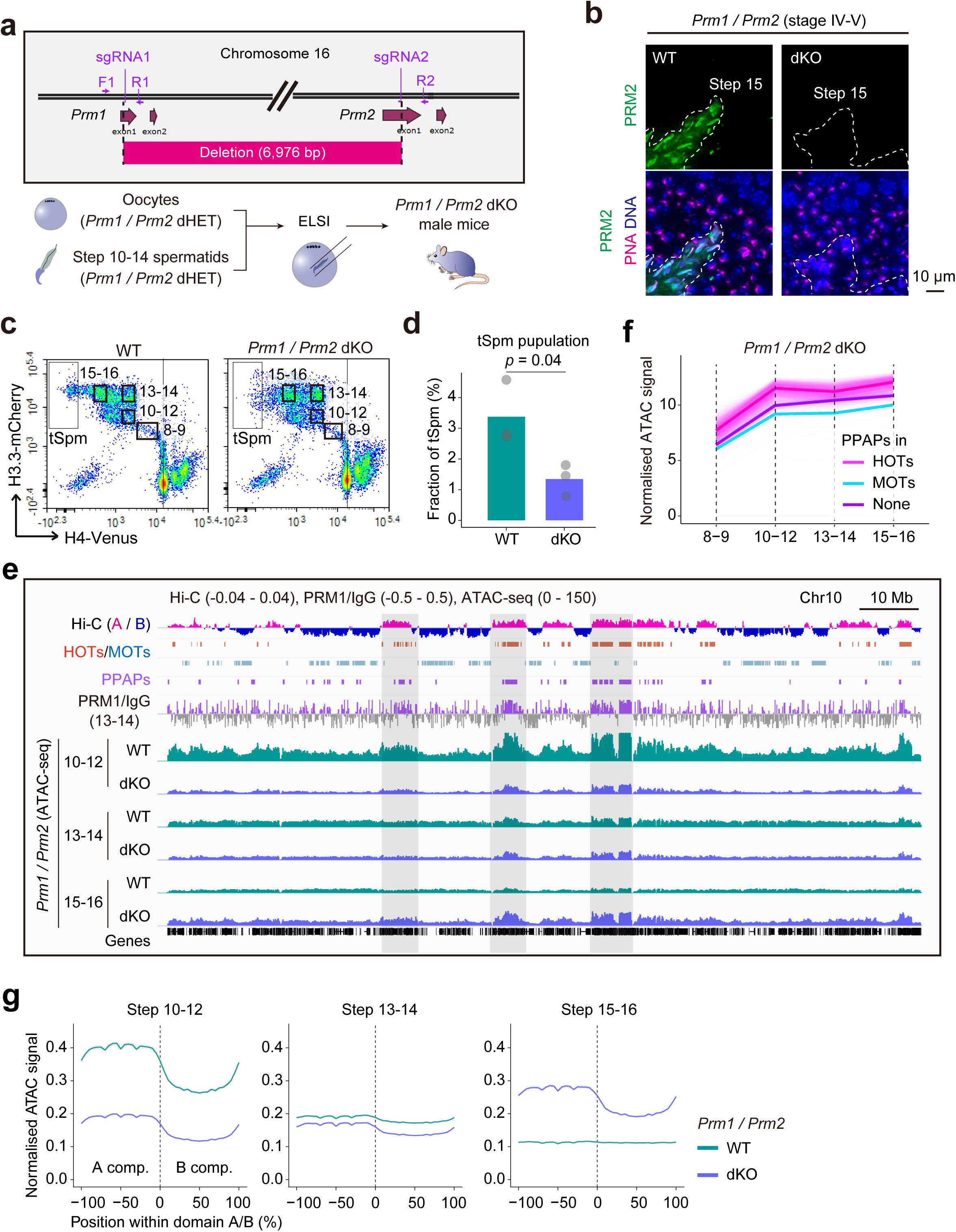
Loss of PRM1 and PRM2 prevents chromatin closure at A compartments during late spermiogenesis. **a**, Schematic illustration of the generation of *Prm1/Prm2* double-knockout (dKO) mice. Guide RNAs targeting the *Prm1* and *Prm2* loci on chromosome 16 were used to generate a 6,976-bp deletion encompassing coding exons. *Prm1/Prm2* dKO male mice were generated by elongated spermatid injection (ELSI) using *Prm1/Prm2* double-heterozygous (dHET) oocytes and spermatids. **b,** Immunofluorescence analysis of PRM2 from WT and *Prm1/Prm2* dKO testes. PRM2 signal was absent in step 15 dKO spermatids. Scale bar, 10 µm. **c,** Cell sorting plots and gate settings showing spermatid populations isolated from WT and *Prm1/Prm2* dKO testes. Populations corresponding steps 8-9, 10-12, 13-14, 15-16, and tSpm populations are indicated. **d,** Quantification of tSpm frequency in WT and *Prm1/Prm2* dKO testes. *p*-value was calculated using a two-sided statistical test. **e,** Genome browser snapshots showing Hi-C PC1 profiles, PRM1 CUT&Tag signal (normalized to IgG), and spike-in–normalized ATAC-seq signals in WT and *Prm1/Prm2* dKO spermatids across late spermiogenic stages. ATAC-seq signals at A compartments remain elevated in *Prm1/Prm2* dKO spermatids at steps 15–16, in contrast to WT spermatids. **f**, Stage-resolved dynamics of spike-in–normalized ATAC-seq signals within PPAPs, HOTs, and MOTs in dKO spermatids. **g**, Average ATAC-seq signal profiles across A/B compartment boundaries in WT and dKO spermatids.

To stage dKO spermatids, we introduced the H4–Venus/H3.3–mCherry transgenes and performed flow-cytometry–based sorting. Although the reporter patterns were broadly preserved compared with *Phf7^CA/CA^* mice (Fig. 3a), the distribution of populations was altered particularly with fewer tSpm (Fig. 5c, d, S5d). ATAC-seq showed that the steps 10–12 accessibility increase was still detectable in dKO spermatids, particularly within A-compartment regions, albeit with reduced amplitude compared with wild type (Fig. 5e). Accessibility then decreased at steps 13–14. However, in steps 15–16 spermatids, only dKO samples showed regained accessibility within A-compartment regions, whereas B-compartment regions were comparatively unchanged (Fig. 5f, 5g). Accessibility of HOT and MOT across spermiogenic stages in dKO exhibited distinct pattern from wildtype; their accessibility kept increased during late spermiogenesis (Fig. 1h, 5f, 5g).

Compartment-stratified differences in accessibility further demonstrated that the dKO showed overall reduced accessibility compared with the wildtype in steps 10-12. In wildtype, the difference between the A- and B-compartments had already diminished by steps 13-14 and became globally inaccessible by steps 15-16, while the dKO exhibited global increase in accessibility at steps 15-16 with a particular increase in accessibility to A, leading to the restoration of A/B compartmentalisation (Fig. 5g). Collectively, observations in *Prm1/Prm2* dKO spermatids suggest that, while the replacement-associated accessibility increase can still occur, PRMs are required to stabilise and maintain late-stage compaction of A-compartment chromatin, preventing aberrant re-opening and the re-emergence of compartment-stratified accessibility.

### DNA fragmentation is initiated at A-compartment region in the absence of PRMs

Epididymal sperm from protamine-deficient models are reported to exhibit extensive DNA damage and fragmentation. In *Prm1* KO males, epididymal sperm DNA is strongly degraded into short fragments (∼100–500 bp) and oxidative lesions are already detectable in caput sperm, whereas in *Prm2* KO males oxidative DNA damage emerges after sperm leave the testis and intensifies during epididymal maturation towards the cauda^21,22^. Based on these studies and our PRM1 maps, we hypothesised that, upon impaired protamination, DNA fragmentation is initiated preferentially within A-compartment regions and subsequently spreads genome-wide during epididymal transit. To test this, sperm were collected from the caput, corpus and cauda epididymis and fragmented DNA from each region was subjected to genomic sequencing. Because DNA from *Prm1/Prm2* dKO cauda sperm was already extensively fragmented, limiting our ability to infer early spatial biases, we performed this analysis in *Prm1/Prm2* dHET males.

Epididymal sperm were identified by retained H3.3–mCherry fluorescence and purified by flow-cytometry–based sorting to exclude contaminating non-sperm cells (Fig. 6a, 6b, S6a-d). In wild-type sperm, H4-Venus intensity decreased from caput to cauda, consistent with progressive chromatin remodelling and/or fluorescence quenching in increasingly compact nuclei (Fig. 6b, c, upper panels). In contrast, *Prm1/Prm2* dHET sperm exhibited recurrent H4-Venus expression in cauda epididymis, implying compromised transcription or auto-fluorescence in defective, moribund sperm (Fig. 6b, c, lower panels). H3.3-mCherry was also slightly decreased in the dHET sperm compared with the wild type (Fig. 6c). DNA extracted from sorted dHET sperm comprised both short (∼200 bp) and high-molecular-weight (>10 kb) fragments in the cauda, whereas the short-fragment fraction became progressively more dominant during epididymal transit (Fig. 6d, arrows), suggesting the propagation of DNA damage. Progressive DNA fragmentation during the epididymal transit in dHET was also confirmed by sperm-optimised comet assay (Fig. S6e, S6f)^23^.

**Figure 6.**
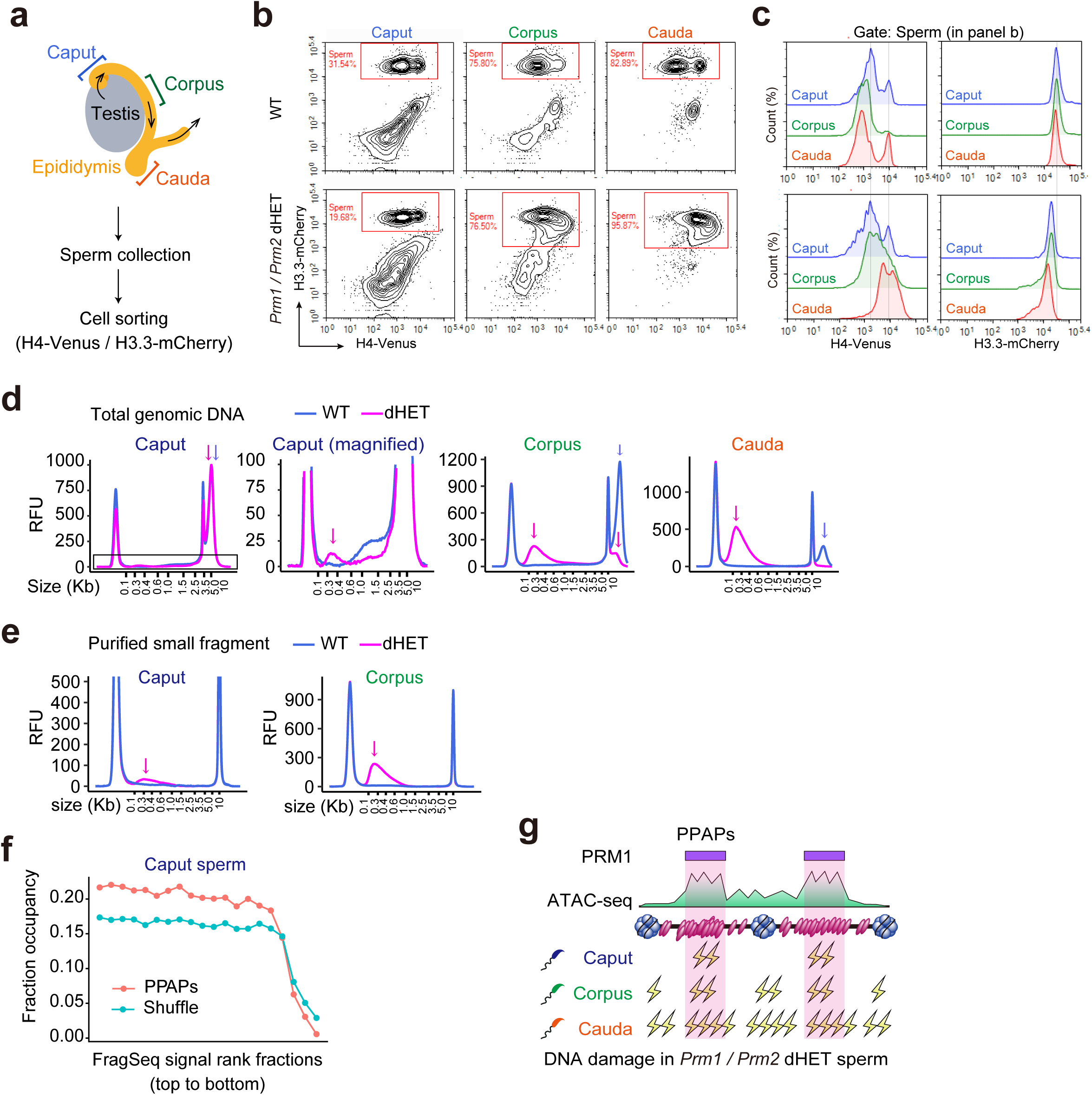
Preferential incorporation of PRM1 into A-compartments protects the sperm genome from DNA damage. **a**, Schematic illustration of the experimental workflow for detecting genome-wide DNA damage in epididymal sperm. Spermatozoa were collected from the caput, corpus, and cauda regions of the epididymis from *Prm1/Prm2* dHET males, followed by isolation of genomic DNA, recovery of low-molecular-weight DNA fragments, and next-generation sequencing. **b,** Cell sorting plots showing sperm populations collected from the caput, corpus, and cauda epididymis of WT and *Prm1/Prm2* dHET males using H4-Venus and H3.3-mCherry reporters. **c,** Flow cytometry histograms showing fluorescence intensity distributions of H4-Venus and H3.3-mCherry signals in epididymal sperm populations. **d,** Size distributions of total genomic DNA isolated from caput, corpus, and cauda sperm from WT and *Prm1/Prm2* dHET males. **e,** Size distributions of purified low-molecular-weight DNA fragments recovered from caput and corpus sperm. **f,** Enrichment of PPAPs among genomic bins ranked by the DNA fragment sequencing signal intensity in the caput sperm. Genomic bins were ordered from highest to lowest sequencing signal, and the fraction overlapping PPAPs was calculated for each rank fraction. **g,** Schematic model illustrating the relationship between PRM1 incorporation, chromatin accessibility, and DNA damage in epididymal sperm. In dHET sperm, DNA damage in cauda sperm is broadly distributed across the genome, whereas caput sperm exhibits preferential DNA damage accumulation within A compartments.

The short-fragment DNA recovered from caput of dHET sperm was purified and sequenced (Fig. 6e, arrows). Importantly, the caput-derived short fragments were enriched at PPAPs, whereas this enrichment was not observed in shuffled control regions, in which PPAPs were randomly redistributed across the mouse genome (Fig. 6f). These results support a model in which fragmentation is initially biased towards protamine-targeted, A-compartment regions and becomes progressively less spatially restricted during epididymal transit when protamination is compromised (Fig. 6g).

As summarised in Fig. 7, our step-resolved ATAC-seq and PRM1 CUT&Tag analyses define a non-monotonic remodelling programme in which a transient, genome-wide chromatin opening accompanies H–P replacement. PRM1 signal first becomes detectable during this window as dispersed, kilobase-scale peaks with only modest compartmental preference, and is subsequently organised into megabase-scale PPAPs enriched in the A compartment before becoming more widespread at later stages. When protamine loading is impaired, DNA fragmentation is initially enriched at protamine-targeted regions during early epididymal transit and progressively spreads genome-wide.

**Figure 7.**
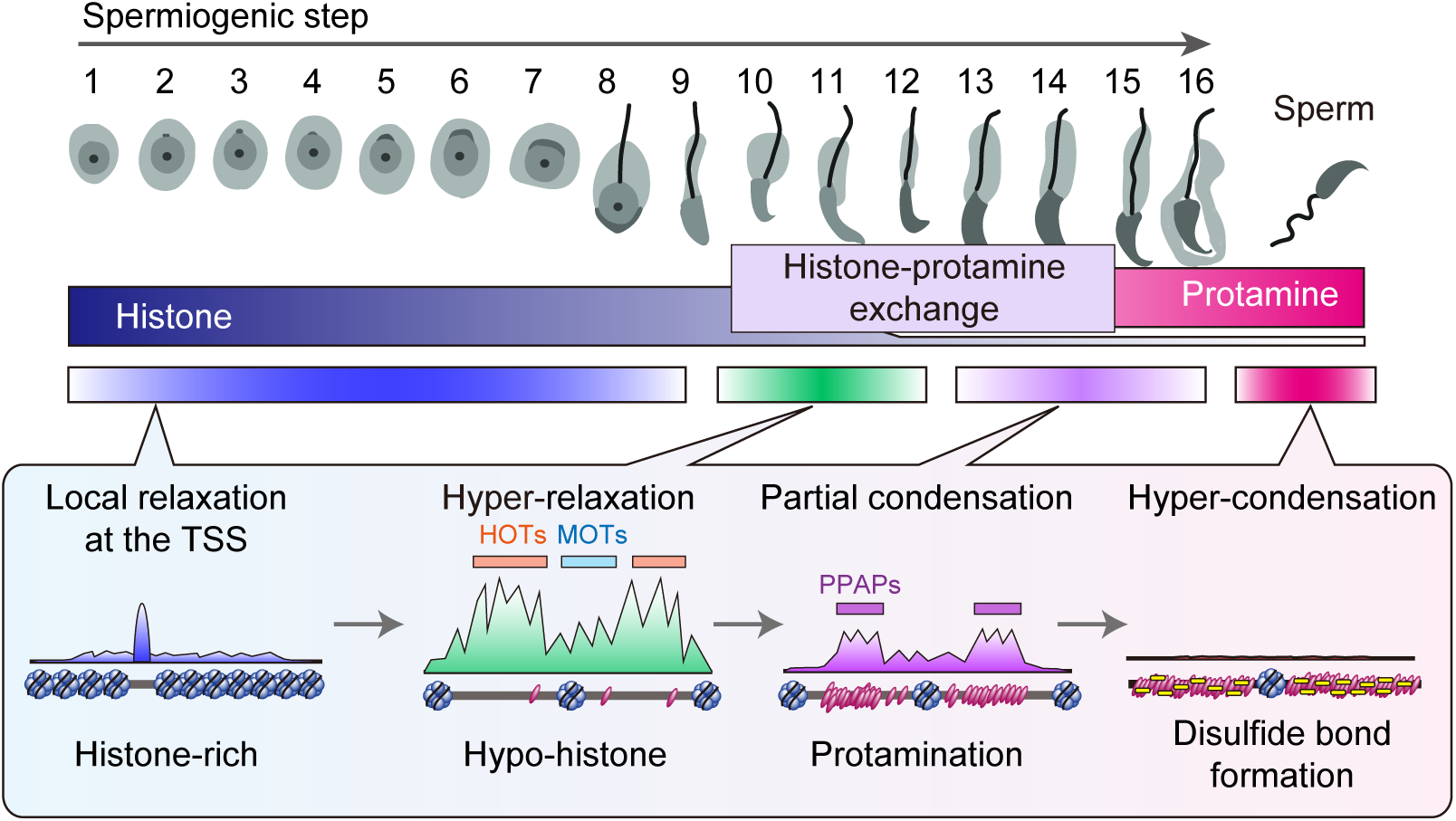
Model for stepwise chromatin remodeling and protamine incorporation during spermiogenesis. Schematic model summarizing step-resolved chromatin remodeling during spermiogenesis. Early spermatids (steps 1–8) retain histone-rich chromatin with localized accessibility at transcription start sites. During mid-spermiogenesis (steps 10–12), chromatin undergoes a transient genome-wide hyper-relaxed state characterized by extensive histone eviction and the emergence of HOTs, predominantly within A compartments. These hyper-accessible regions constitute PPAPs and serve as major sites of PRM1 incorporation during late spermiogenesis (steps 13–14). Protamine deposition is coupled to rapid chromatin closing, resulting in progressive compartment-biased condensation. In the absence of protamines, hyper-accessible A-compartment regions fail to undergo proper closure, leading to persistent accessibility and preferential DNA fragmentation. This non-monotonic remodeling trajectory establishes a spatially organized transition from histone-based to protamine-based chromatin architecture during sperm maturation.

## Discussion

Early genome-wide surveys of sperm chromatin—spanning nuclease sensitivity, nucleosome mapping and epigenomic profiling—revealed pronounced contrasts between histone- and protamine-dominated packaging and implied extensive remodelling during spermiogenesis^24–28^. In particular, ATAC-seq–based studies tied waves of H4 hyperacetylation and BET-reader activity to nucleosome dynamics and histone eviction^29^, but left unresolved how the moment of replacement is organised in vivo. Here, by using highly purified spermatids isolated from each spermiogenic step and combining spike-in-normalised ATAC-seq with CUT&Tag, we identify a previously unappreciated transient, genome-wide hyper-relaxation that bridges histone eviction and protamine loading, and we further link this transition to the compartmental structure of the genome.

A first conclusion is that histone–protamine replacement is neither monotonic in time nor uniform across the genome. Spermatids traverse a hyper-relaxed state (steps 10–12) in which nucleosomal phasing is attenuated and accessibility increases broadly, with the strongest changes in A-compartment, gene-rich regions, before accessibility subsequently declines towards terminal compaction. Notably, this transient loosening is not coupled with transcriptional output (Fig. S2). Conceptually, these data refine earlier observations that chromatin accessibility changes across spermatogenesis by pinpointing a brief, genome-wide opening immediately prior to terminal compaction that was not resolved by conventional coarse fractionation-based analyses of spermatids^29^.

Mechanistically, our PHF7 experiments place histone eviction upstream of this opening. In a catalytic PHF7 mutant, the hyper-relaxed state is suppressed and nucleosomal ladders persist, even though PRM1/PRM2 levels are comparatively maintained and H4 acetylation is increased^7^. This genetic separation indicates that protamine availability per se is insufficient: effective sperm chromatin compaction and genome protection require an eviction-licensed, transiently open, largely histone-depleted substrate that permits productive protamination and higher-order packaging. In addition, previous reports that PHF7 localises preferentially to gene-rich regions are consistent with our observation that accessibility increases most strongly within A-compartment domains^6^.

A second conclusion is that initiation of protamine loading is spatially selective. PRM1 CUT&Tag mapping indicates preferential incorporation within the A compartment, and local PRM1 enrichment correlates with underlying chromatin accessibility (Fig. 4b). This may reflect preferential deposition into transiently accessible chromatin, and/or additional biases linked to A-compartment identity. For example, biophysical and computational studies suggest that DNA base composition and groove geometry can modulate protamine–DNA interaction modes and the efficiency of protamine-driven collapse, with GC-rich templates favouring tighter condensation under some conditions^30^. Because A-compartments tend to be gene-dense and relatively GC-rich, such intrinsic DNA properties together with higher accessibility could contribute to preferential PRM1 loading into A-compartment chromatin. Importantly, this spatial bias also has functional consequences: in protamine-deficient settings, fragmented DNA is enriched at protamine-targeted regions early during epididymal transit and becomes progressively less spatially restricted as fragmentation spreads genome-wide, linking compartment-biased protamine deposition to compartment-biased vulnerability.

How, then, is compaction consolidated across B-compartment chromatin, and is delayed protamine deposition compatible with genome protection? Although PRM1 becomes detectable during steps 10–12 as dispersed, kilobase-scale enrichment with only modest compartmental preference, it is subsequently organised in steps 13–14 into A-enriched PPAPs before spreading more broadly into both compartments at later stages. This suggests that B domains are not exempt from protamination, but may undergo later consolidation relative to A-compartment. Consistent with such a temporal hierarchy, in protamine-deficient settings fragmentation is initially enriched at protamine-targeted regions and then progressively loses spatial restriction as damage spreads genome-wide during epididymal transit. In this case, how might peripheral B chromatin tolerate delayed protamine consolidation? In canonical A/B compartmentalisation, B domains correspond to transcriptionally inert, heterochromatin-like chromatin and are frequently positioned at the nuclear periphery, where additional physical constraints could provide baseline compaction^31–33^. In somatic nuclei, transcriptionally-suppressed chromatin often overlaps lamina-associated domains tethered to the nuclear lamina, and spermiogenesis entails extensive nuclear-envelope remodelling^34,35^; whether analogous perinuclear mechanisms help buffer B-compartment compaction and integrity before full protamination is completed remains an important open question.

In parallel, the persistence of nuclear shrinkage in *Prm1/Prm2*-deficient spermatids suggests that transition proteins and other basic chromatin factors can further buffer compaction when protamines are limiting. One non-mutually exclusive buffer is compensation by transition nuclear proteins. TNP1/TNP2 normally occupy chromatin transiently during elongation and promote an intermediate compaction state before being replaced by protamines; consistent with a dosage-sensitive buffering of this histone→TNP→protamine sequence, disruption of *Tnp1* or *Tnp2* perturbs condensation and is accompanied by compensatory changes in other basic nuclear proteins^36–39^. In our *Prm1/Prm2* dKO spermatids, increased TNP1 abundance (Fig. S5b) raises the possibility that prolonged transition-protein occupancy contributes to residual nuclear shrinkage when protamines are limiting.

Bringing these strands together, our data motivate a domain-aware model of paternal genome packaging. Gene-rich, transcription-competent A compartments undergo PHF7-licensed, eviction-triggered hyper-relaxation, followed by early protamine loading that stabilises A-compartment closure. Protamine deposition then becomes more widespread, extending into B-compartment chromatin to consolidate terminal genome-wide compaction, whereas baseline compaction of gene-poor, peripherally positioned B domains may be buffered before full protamination by transition proteins and/or perinuclear constraints. In addition, although not directly addressed in this study, distinct condensation mechanisms are likely applied at centromeres, where specialized nucleosome-based chromatin (including CENP-A and phosphorylated TH2A) is retained, and on sex chromosomes, which are tightly packaged into heterochromatin^40–42^. Together, these layers suggest that sperm genome protection emerges from partially parallel, context-dependent mechanisms rather than from a single uniform condensation pathway.

Collectively, our results reposition histone–protamine replacement as a multi-step, domain-aware programme: PHF7-dependent eviction licenses a transient genome-wide opening, and protamination preferentially stabilises compaction within relaxed A-compartment chromatin, while additional protamine-independent forces might contribute elsewhere.

## Materials and methods

### Mouse and ethics statement

C57BL/6 mice (Japan SLC) were sacrificed by cervical dislocation and used for experiments. All animal care and experimental procedures were approved by the Animal Experiment Ethics Committees at the Institute for Quantitative Biosciences, The University of Tokyo (approval numbers 0212, 0313, and 0409), and were performed in strict accordance with the guidelines provided by the Life Science Research Ethics and Safety Office of The University of Tokyo. Homozygous H4-Venus/H3.3-mCherry (VC) transgenic mice^18,43^ were maintained on a C57BL/6J background. *Phf7* C160A mutant (CA) mice^7^ were crossed with VC mice.

### Antibodies, sgRNAs and primers

Antibodies used in this study were listed in Supplemental Table S1. Information of sgRNAs for CRISPR/Cas9 and primers were provided in Supplemental Table S2.

### Flow cytometry-based cell sorting of spermatids

Flow cytometry-based cell isolation of spermatids was performed as previously described with minor modifications^18^. Briefly, the tunica albuginea was removed from adult mouse testes, which were then gently teased apart with tweezers and transferred to PBS containing 1.0 mg/mL collagenase type I (Sigma-Aldrich). Tissues were incubated at 37°C for 5–10 min until the seminiferous tubules were dispersed. After centrifugation, the supernatant was discarded and 0.5% trypsin (Thermo Fisher) containing 2 mg/mL DNase I (Sigma-Aldrich) was added to the pellet. The pellet was briefly pipetted up and down and incubated at 37°C for 5 min. An equal volume of fetal bovine serum (FBS) was then added to inactivate trypsin, and the cell/FBS mixture was pipetted up and down and left at room temperature for 3 min. Cells were filtered through a 70-µm cell strainer and washed three times with PBS containing 2% FBS (PBS–FBS). For DNA content and viability staining, cells were resuspended in DMEM (Wako) containing 10% FBS and 5 µM Vybrant DyeCycle Violet (DCV; Invitrogen) and incubated at 37°C for 30 min. Immediately before sorting, DRAQ7 (BioStatus) was added at a final concentration of 3 mM and the cells were incubated for 15 min at room temperature. Cells were then subjected to cell sorting.

Cell sorting was performed on a BD FACSAria III using BD FACSDiva software. A 100-µm nozzle was used. Excitation was provided by 405-, 488-, 561-, and 633-nm lasers. The sort mode was set to 4-Way Purity with a flow rate of 1.0–3.0, resulting in a processing rate of ∼5,000 events per second. At this rate, bilateral testes from one adult mouse could be processed within 6–7 h. Sorted cells were collected into 5-mL flow cytometry tubes containing 0.5% BSA/PBS. The filter sets used for sorting and detection were as follows: (i) DAPI/DCV: 450/40, (ii) Venus: 502LP/530/30, (iii) mCherry: 600LP/610/20, and (iv) DRAQ7: 660/20. DRAQ7-negative, DCV-positive populations were collected as healthy testicular cell population. The detailed gate settings for each spermatid are shown in the Results section. NovoExpress (Agilent Technologies) was used for data analysis and visualization.

### Spike-in ATAC-seq library preparation

Libraries for ATAC-seq were prepared as previously described with minor modifications^44,45^. Sorted spermatids or sperm were washed once with 0.5% BSA in PBS, counted using a hemacytometer, and 10,000 cells were collected per sample. For spike-in normalization, 2,500 Drosophila melanogaster S2 cells were added to each sample. After an additional wash with PBS, cells were resuspended in 50 µL of ice-cold lysis buffer (10 mM Tris-HCl pH 7.5, 10 mM NaCl, 3 mM MgCl₂, 0.1% NP-40, 0.1% Tween-20, 0.01% digitonin), gently pipetted up and down five times, and incubated on ice for 5 min. Nuclei were washed by adding 1 mL of wash buffer (10 mM Tris-HCl pH 7.5, 10 mM NaCl, 3 mM MgCl₂, 0.1% Tween-20), gently inverting the tube, and collecting the nuclear pellet by centrifugation. After removal of the supernatant, the nuclear pellet was resuspended in 50 µL of Tn5 reaction mixture (10 mM Tris-HCl pH 7.4, 5 mM MgCl₂, 10% dimethylformamide, 0.05% digitonin, 0.1% Tween-20, 0.33× PBS, and in-house Tn5 transposase at 100 nM) and incubated at 37°C for 30 min with shaking. Tagmented DNA was purified using a MinElute Reaction Cleanup Kit (Qiagen), and libraries were amplified by PCR using NEBNext High-Fidelity 2× PCR Master Mix (New England Biolabs) and Nextera Ad1/Ad2 index primers. The number of PCR cycles was determined for each sample by quantitative PCR, choosing the cycle number corresponding to one-third of the maximal fluorescence. Final PCR products were purified and size-selected with AMPure XP beads (Beckman Coulter) to enrich for appropriately sized DNA fragments. Libraries were sequenced on an Illumina HiSeq X Ten or NovaSeq platform in 150-bp paired-end mode.

### RNA-seq library preparation for isolated spermatids

Total RNA was isolated from FACS-sorted spermatids using the NucleoSpin RNA XS Plus kit (TaKaRa) according to the manufacturer’s instructions. RNA-seq libraries were generated using the SMART-Seq HT Kit (TaKaRa). Libraries were sequenced on an Illumina HiSeq X Ten platform in 150-bp paired-end mode.

### CUT&Tag library preparation

CUT&Tag was performed essentially as described previously with minor modifications^46^. Briefly, cells were pelleted, resuspended in ice-cold NE1 buffer (20 mM HEPES-KOH pH 7.9, 150 mM NaCl, 0.5 mM spermidine, 1% NP-40, 1% BSA, EDTA-free protease inhibitors), and incubated on ice for 10 min. Nuclei were collected by centrifugation, resuspended in PBS-binding buffer (1× PBS pH 6.8, 1 mM CaCl₂, 1 mM MnCl₂, 1% BSA, protease inhibitors), and lightly fixed with 0.1% formaldehyde for 2 min at room temperature. Cross-linking was quenched by adding glycine to a final concentration of 0.25 M and incubating for 5 min. Nuclei were then incubated with concanavalin A–conjugated Dynabeads MyOne T1 in binding buffer (20 mM HEPES pH 7.5, 10 mM KCl, 1 mM CaCl₂, 1 mM MnCl₂, 1% BSA) for 10 min at room temperature with rotation, washed, and resuspended in NE2 buffer (20 mM HEPES-KOH pH 7.9, 150 mM NaCl, 0.5 mM spermidine, 0.05% Triton X-100, 10% glycerol, 1% BSA, protease inhibitors).

Bead-bound nuclei were washed in wash buffer (20 mM HEPES pH 7.5, 150 mM NaCl, 0.5 mM spermidine, 1% BSA, 1 mM CaCl₂, 1 mM MnCl₂, protease inhibitors) and then incubated in 50 µL antibody buffer (wash buffer containing 0.05% digitonin and 2 mM EDTA) with 0.5 µg primary antibody overnight at 4°C with gentle mixing. After washing in Dig-wash buffer (wash buffer with 0.05% digitonin), cells were incubated for 1 h at room temperature with secondary antibody (1:100; ROCKLAND 610-4120) in Dig-wash buffer, washed 2–3 times, and then incubated for 1 h at room temperature in Dig-300 buffer (20 mM HEPES pH 7.5, 300 mM NaCl, 0.5 mM spermidine, 0.01% digitonin, protease inhibitors) containing 50 nM pA–Tn5 transposome. After two washes with Dig-300 buffer, tagmentation was performed by resuspending the beads in 200 µL tagmentation buffer (Dig-300 buffer with 10 mM MgCl₂) and incubating at 37°C for 1 h. Reactions were stopped by adding 0.5 M EDTA, 10% SDS, and proteinase K, followed by incubation at 56°C for 3 h or overnight. DNA was purified using a MinElute Reaction Cleanup Kit (Qiagen), and CUT&Tag libraries were generated by PCR using NEBNext High-Fidelity 2× PCR Master Mix and indexed primers. The optimal number of PCR cycles was determined by a side quantitative PCR reaction, after which the remaining library was amplified, size-selected with AMPure XP beads, and sequenced on a DNBSEQ-G400 in 150-bp paired-end mode.

### Small fragment DNA sequencing from epididymal sperm

Spermatozoa were collected separately from the caput, corpus, and cauda regions of the epididymis. For caput and corpus samples, dissected epididymal tissues were placed in HTF medium and mechanically dissociated using fine forceps to release sperm. The resulting cell suspensions were passed through a 35-µm cell strainer and subjected to cell sorting to purify spermatozoa based on H3.3-mCherry signal. Genomic DNA was extracted from the sorted sperm using sperm genome extraction buffer (50 mM Tris-HCl [pH 8.0], 10 mM EDTA, 0.5 M NaCl, 0.2% SDS, 10 mM DTT, and 0.25 mg/mL proteinase K), followed by phenol–chloroform–isoamyl alcohol extraction and ethanol precipitation. Fragment size distributions of the extracted genomic DNA were assessed using a TapeStation system with the High Sensitivity D5000 kit. For caput and corpus samples, low-molecular-weight DNA fragments were selectively enriched from total genomic DNA using AMPure bead-based size selection. Purified DNA fragments were used to construct sequencing libraries using the NEBNext Ultra II DNA Library Prep Kit. Libraries were sequenced on an Illumina NextSeq 2000 platform in 36-bp paired-end mode.

### Spike-in–normalized ATAC-seq data processing

Paired-end ATAC-seq reads were first subjected to adapter and quality trimming and then aligned to a combined reference genome consisting of mouse (mm39) and Drosophila melanogaster (dm6) using Bowtie2 with the following parameters: -N 1 -X 2000 --no-mixed --no-discordant. Aligned reads were converted to BAM format and coordinate-sorted using Samtools. Mitochondrial reads (chrM) were removed, and only properly paired reads with a mapping quality ≥30 were retained. PCR duplicates were removed using the collate–fixmate–sort–markdup workflow in Samtools. The resulting BAM files were split into mouse and Drosophila alignments based on the reference genome, and reads overlapping blacklist regions in either genome were removed using bedtools and pre-defined mm39 and dm6 blacklist BED files. For each sample, the numbers of blacklist-removed reads in mouse and Drosophila genomes were summarized, and these counts were used to calculate sample-specific spike-in normalization factors for downstream analyses.

### CUT&Tag data processing

Adapter and low-quality bases were trimmed using Trim Galore in paired-end mode. Trimmed reads were aligned to the mouse reference genome (mm39) using Bowtie2 with the following parameters: *-N 1 -X 2000 --no-mixed --no-discordant*. Alignments were converted to BAM format, coordinate-sorted, and indexed using Samtools. Subsequently, alignments were filtered to retain high-confidence mapped reads (MAPQ ≥30). Reads mapping to the mitochondrial chromosome (chrM) were removed. PCR duplicates were removed from paired-end alignments using a Samtools-based duplicate marking/removal workflow. In addition, reads overlapping the ENCODE mm39 blacklist were excluded. These filtered BAM files were used for replicate concordance assessment. Genome-wide similarity between biological replicates was evaluated by computing Pearson correlation coefficients from binned genome-wide coverage profiles. Replicates with Pearson correlation coefficients ≥0.7 were merged and the merged datasets were used for downstream analyses.

### Generation of Prm1 and Prm2 double-knockout mice

MII-stage oocytes were collected from 8–12-week-old super-ovulated VC female mice. Superovulation was induced by intraperitoneal injection of a mixture of 2.5 IU pregnant mare serum gonadotropin (PMSG; ASKA Pharmaceutical) and 100 μL anti-inhibin serum (Central Research), followed 48 h later by 7.5 IU human chorionic gonadotropin (hCG; ASKA Pharmaceutical). For in vitro fertilization, MII oocytes were inseminated with capacitated spermatozoa obtained from the cauda epididymides of 10-week-old VC male mice. Zygotes were transferred into M2 medium and subjected to microinjection. Approximately 2–4 pL of injection mixture containing 50 ng/μL Cas9 mRNA (TriLink Biotechnologies) and 0.4 pM each of guide RNA1 and guide RNA2 targeting *Prm1* and *Prm2* genome (Synthego) was injected into the cytoplasm of zygotes using a pneumatic microinjector (Narishige) equipped with a piezo micromanipulator (Prime Tech). After injection, zygotes were cultured overnight in HTF medium. Two-cell embryos were then transferred into the uteri of pseudopregnant female mice. Resulting *Prm1/Prm2* dHET females were backcrossed with VC males for at least three generations to dilute potential off-target effects. *Prm1/Prm2* dKO male mice were subsequently generated by injecting condensed spermatids from dHET males into oocytes from dHET females using a piezo-assisted microinjection system. The resulting two-cell embryos were transferred into the uteri of pseudopregnant females.

### Preparation of frozen testis sections and immunohistochemistry

Dissected testes were embedded into optimal cutting temperature (OCT) compound (Sakura Finetek), then sliced sections were fixed with 1% PFA/0.1% Triton X-100. These sections were treated with blocking buffer (5 % goat serum (Vector) in TBST) followed by immunostaining. After blocking, sections were incubated with primary antibodies diluted with blocking buffer, and then incubated with Alexa488-, 568-, 647-conjugated secondary antibodies (Life) and Hoechst 33342 (Invitrogen). Images were acquired using a FV1000 or 3000 confocal laser scanning microscope (Olympus) or DeltaVision system (GE). Data were processed using FLUOVIEW software (Olympus) and ImageJ software.

## Data availability

All ATAC-seq, CUT&Tag, and RNA-seq data generated in this study have been deposited in NCBI. The data will be made publicly available once this study has undergone peer review and been accepted for publication in another journal.

## Reproducibility

All experiments were performed at least twice to confirm their reproducibility.

## Contributions

M.Ha. and Y.O. conceived the study, performed experiments and wrote the manuscript. C.Z. performed the comet assays related to Fig. S6f. E.I. and Y.F. assisted with mouse generation. Y.F. and C.K. operated FACS. S.S. and H.K. provided materials required for next-generation sequencing experiments. M.Ho. and S.C. contributed to experiments related to Fig. 6. M.I. and S.H.B. provided the Phf7^CA^ mouse line and contributed to experimental design and discussion. All authors reviewed and approved the manuscript.

## Competing interests

The authors declare not competing interests.

## Acknowledgement

We are grateful to R. Nakato for the generous use of his computational resources for bioinformatic analyses, to TeMIC microscope facility for image acquisition, and to IQB mouse reconstitution service for generating genome-edited mice. We are grateful for S. S. Hammoud (University of Michigan Medical School), A. H. F. M. Peters (Friedrich Miescher Institute for Biomedical Research), and O. J. Rando (UMass Chan Medical School) for helpful discussions. Lastly, we thank all members of Y.O.’s laboratory for helpful discussions and support. This research was funded by MEXT KAKENHI (Grant No. 20H05939 to Y. O., Grant No. 23K14191 to M. Ha., Grant No. JP23H05475 and JP24H02328 to H. K.), JST ERATO (Grant No. JPMJER190 to H. K.), JST CREST (Grant No. JPMJCR24T3 to H. K.), AMED BINDS (Grant No. JP25ama121009 to H. K.), the Creative Research Initiatives Program from the National Research Foundation grant funded by the Korea government (2017R1A3B1023387 to S.H.B.),

---

**Supplementary Figure 1.**
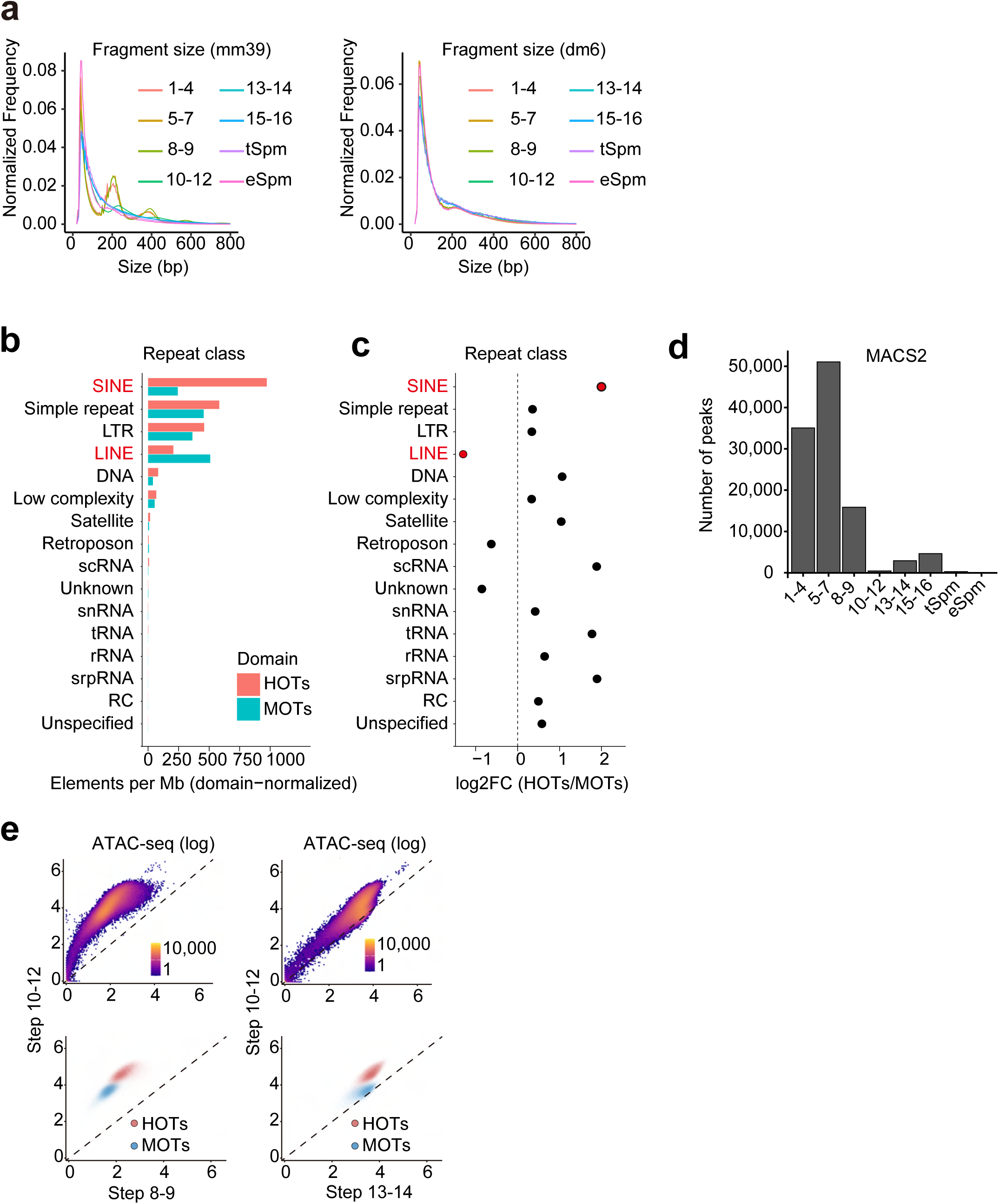
Technical validation and genomic features of chromatin accessibility territories defined by ATAC-seq. **a**, Fragment size distributions of spike-in–normalized ATAC-seq libraries mapped to the mouse genome (mm39; left) and Drosophila genome (dm6; right) across spermiogenic stages and epididymal sperm. **b,** Genomic annotation of HOTs and MOTs based on RepeatMasker classifications. Bar plots show the number of annotated repeat elements per megabase, normalized by domain length. **c,** Relative enrichment of repeat classes in HOTs compared with MOTs, shown as log₂ fold change (HOTs/MOTs). **d,** Distribution of the number of ATAC-seq peaks identified by MACS2 across spermiogenic stages and epididymal sperm. **e,** Scatter plots showing correlations of log₂-transformed ATAC-seq signals between steps 8–9 and steps 10–12 spermatids (left), and between steps 10–12 and steps 13–14 spermatids (right). Regions corresponding to HOTs and MOTs are highlighted.

**Supplementary Figure 2.**
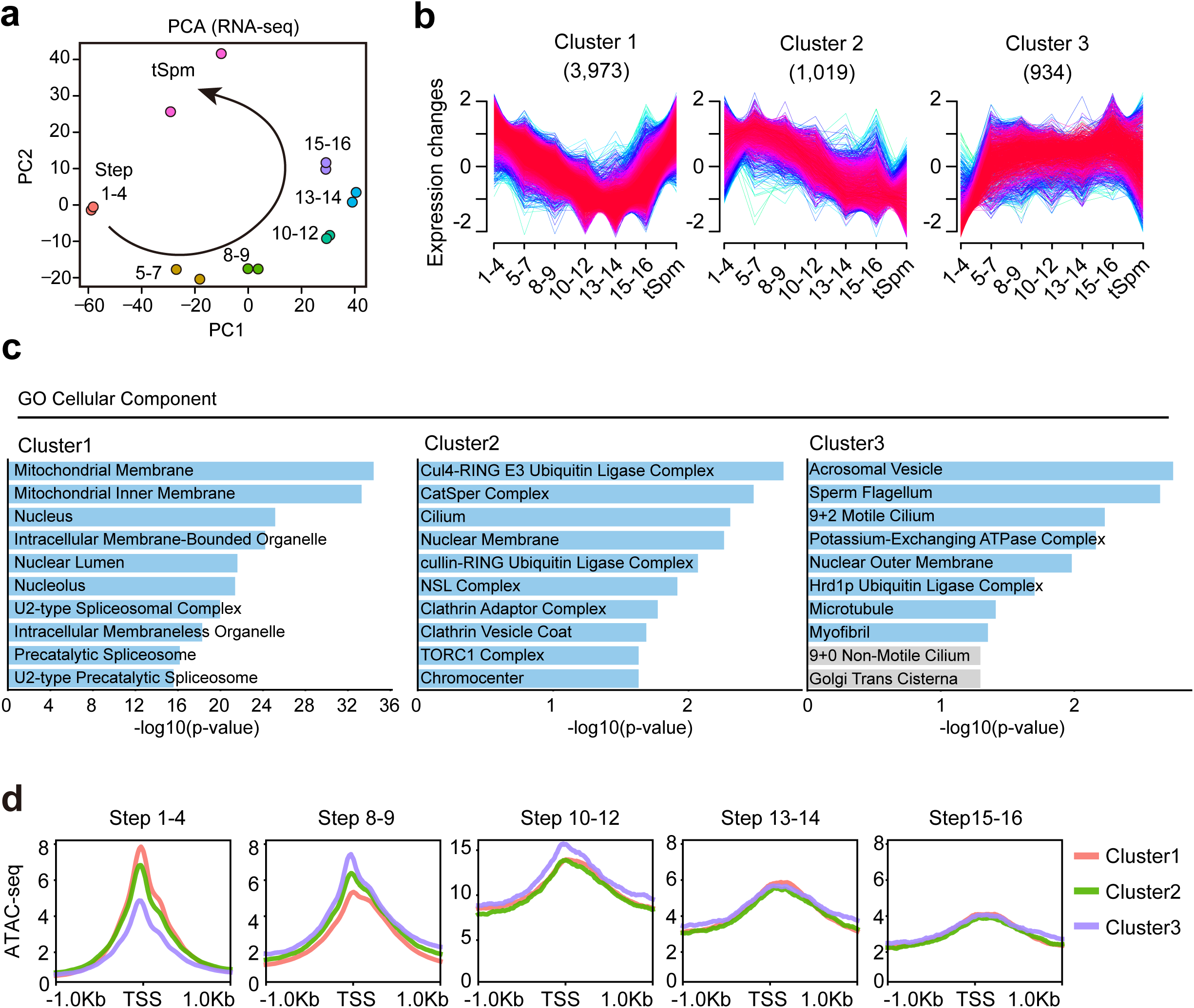
Transcriptome dynamics across spermiogenesis and their relationship to chromatin accessibility. **a**, Principal component analysis (PCA) of RNA-seq data across spermiogenic stages and tSpm. Samples are arranged along a continuous trajectory reflecting progressive spermiogenic differentiation. **b,** Clustering of genes based on their expression dynamics across spermiogenic stages. Line plots show normalized expression changes for individual genes within each cluster, with the number of genes in each cluster indicated. **c,** Gene Ontology (GO) enrichment analysis for Cellular Component terms associated with each expression cluster. Bars indicate −log₁₀-transformed p values for significantly enriched terms. **d,** Average ATAC-seq signal profiles around transcription start sites (TSSs) for genes belonging to each expression cluster, shown across spermiogenic stages. Profiles are centered on the TSS (±1 kb).

**Supplementary Figure 3.**
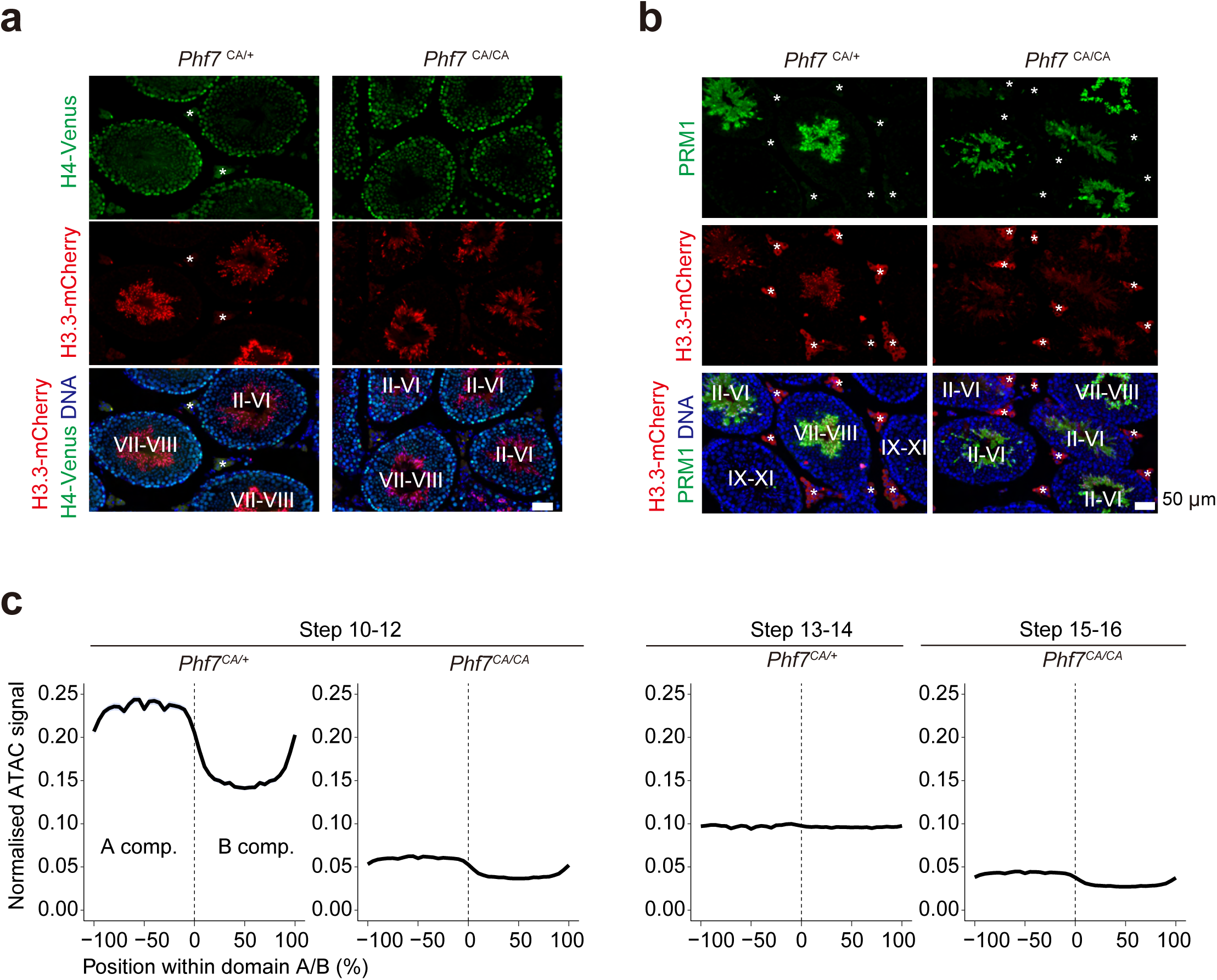
Validation of histone eviction and PRM1 incorporation in Phf7-catalytic mutant spermatids. **a**, Fluorescence images of H3.3-mCherry and H4-Venus signals in seminiferous tubules from control (*Phf7*^CA/+^) and catalytic mutant (*Phf7*^CA/CA^) testes. Spermiogenic stages are indicated in merged images. Asterisks indicate regions showing non-specific fluorescence signals. **b,** Immunofluorescence analysis of PRM1 and H3.3-mCherry signals in seminiferous tubules from *Phf7*^CA/+^ and *Phf7*^CA/CA^ testes. Roman numbers indicated the spermatogenic stages of seminiferous tubules. Asterisks indicate regions showing non-specific fluorescence signals in the interstitial regions. Scale bar, 50 µm. **c,** Average ATAC-seq signal profiles across A/B compartment boundaries in *Phf7*^CA/+^ and *Phf7*^CA/CA^ spermatids. Relative ATAC-seq signal is plotted as a function of genomic position within A and B compartments, illustrating impaired chromatin remodeling in the catalytic mutant.

**Supplementary Figure 4.**
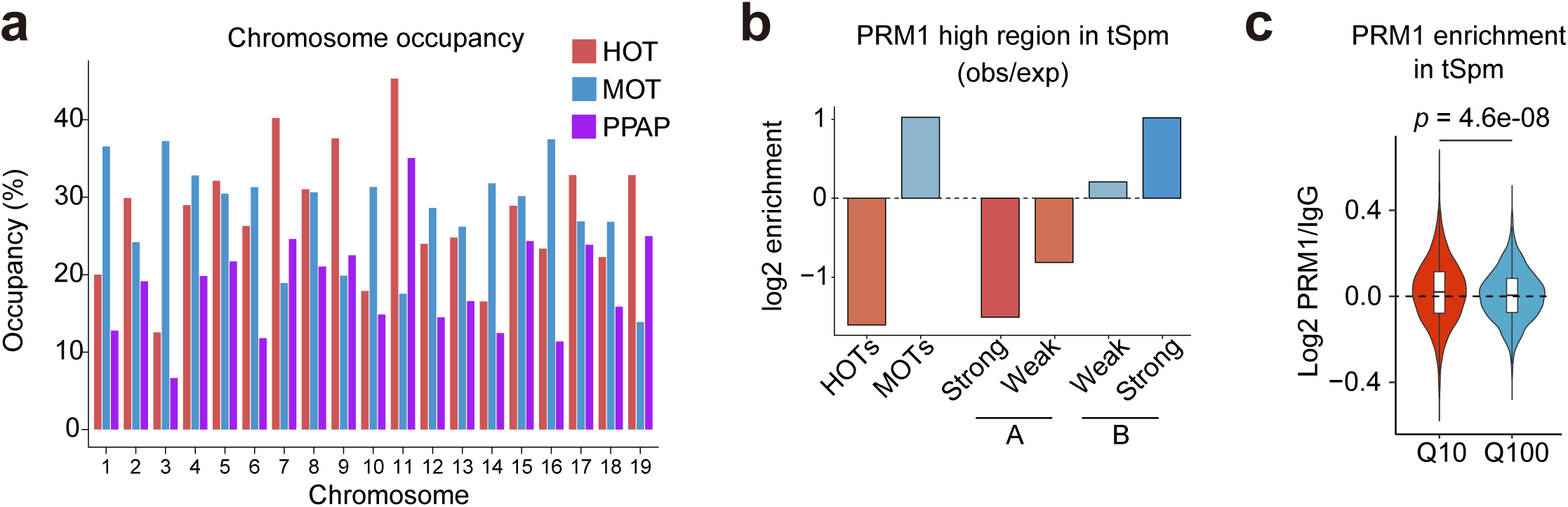
Genomic distribution and enrichment of PRM1-associated regions in testicular sperm. **a**, Chromosome-wide occupancy of HOTs, MOTs, and PPAPs. Bars indicate the percentage of genomic bins assigned to each category across chromosomes 1–19. **b**, Observed/expected (obs/exp) enrichment of PRM1-high regions in tSpm across HOTs, MOTs, and A/B compartment subclasses (strong A, weak A, strong B, weak B). Values are shown as log₂ enrichment. **c**, PRM1 CUT&Tag enrichment (log₂ PRM1/IgG) in tSpm stratified by percentile groups (Q10 and Q100). *P-*value was calculated using a two-sided Welch’s t-test.

**Supplementary Figure 5.**
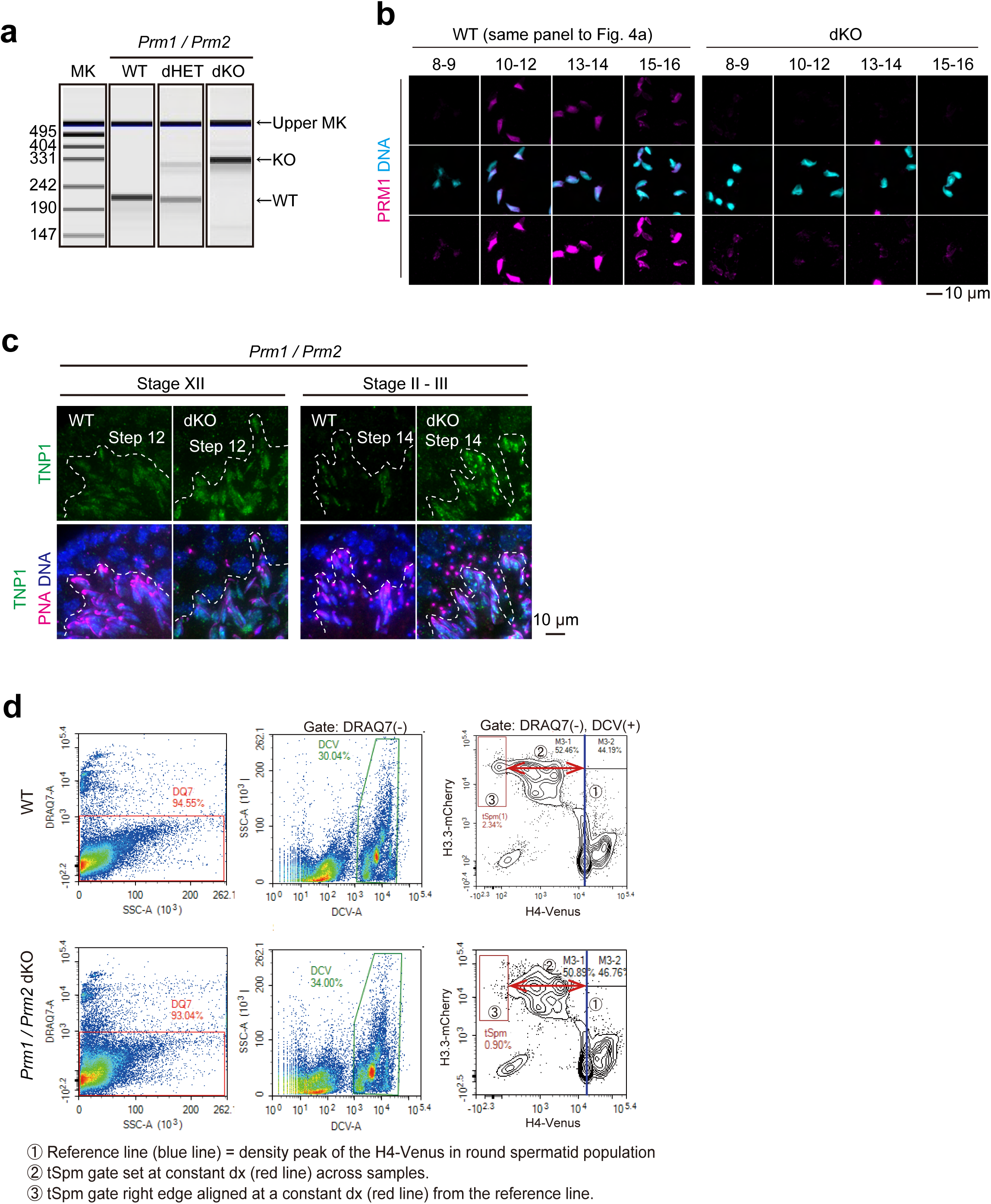
Validation of protamine loss and spermatid phenotypes in Prm1/Prm2 mutant mice. **a**, Genotyping analysis of *Prm1/Prm2* wild type (WT), dounle heterozygous (dHET), and double knockout (dKO mice). Representative electrophoresis images show diagnostic bands corresponding to WT (amplified with F1-R1 primer set in Fig. 5) and KO (amplified with F1-R2 primer set in Fig. 5) alleles, respectively. **b,** Immunofluorescence analysis of PRM1 (magenta) and DNA (cyan) in purified WT and dKO spermatids. Scale bar, 10 μm. **c** Immunofluorescence analysis of TNP1 in seminiferous tubules from WT and *Prm1/Prm2* dKO testes. Representative images are shown for stage XII (step 12) and stages II–III (step 14). PNA and DNA counterstaining delineate acrosome morphology and nuclear structure. Scale bar, 10 µm. **d,** Flow cytometry–based gating strategy for quantification of tSpm populations.

**Supplementary Figure 6.**
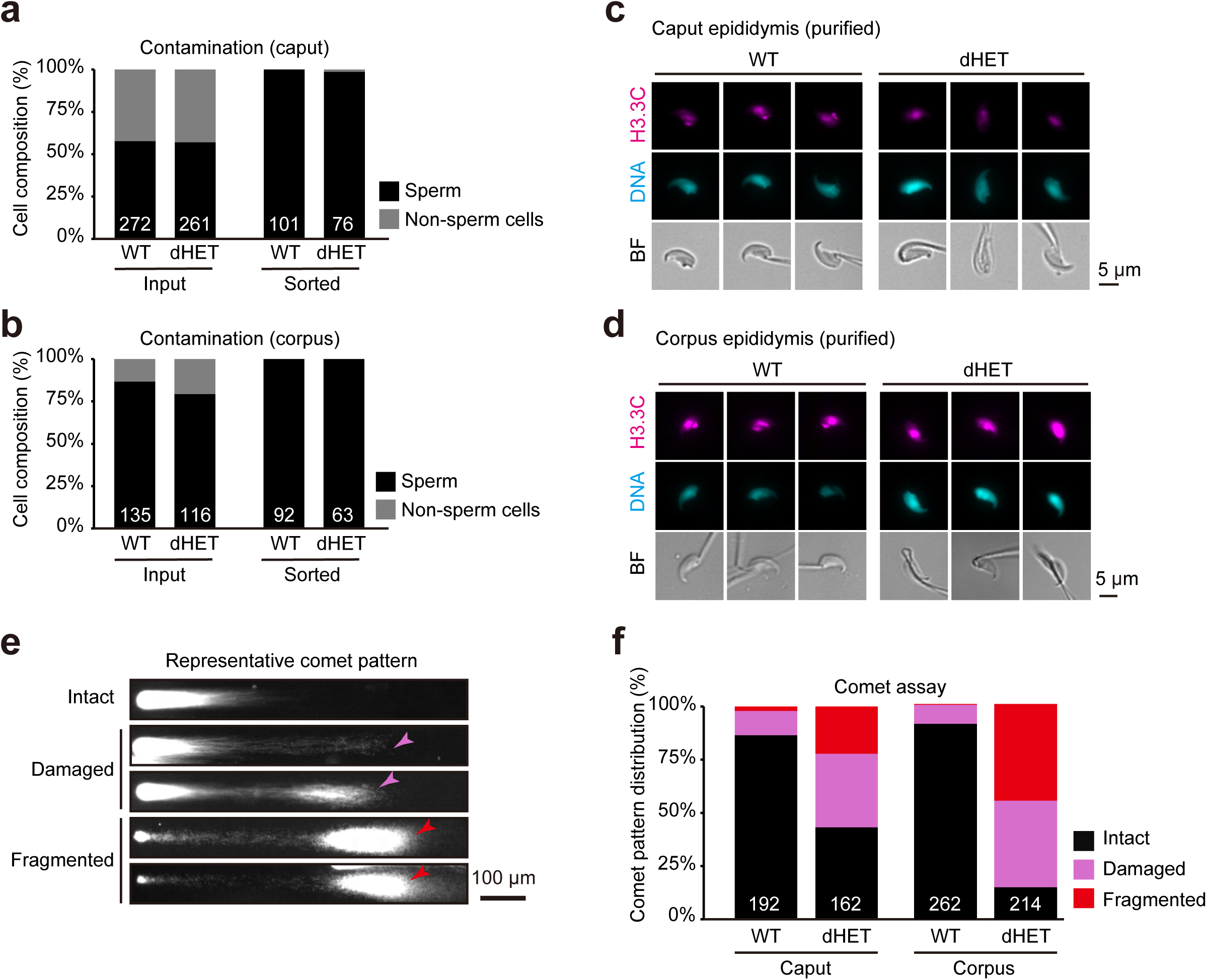
Quality control and validation of DNA damage analyses in epididymal sperm. **a, b,** Quantification of cellular composition in caput (**a**) and corpus (**b**) epididymal samples before (input) and after H3.3-mCherry(+) cell sorting (also see Fig. 6a, 6b). Stacked bar plots show the proportions of sperm and non-sperm cells in WT and dHET samples. Numbers below bars indicate total counted cells. **c, d,** Representative fluorescence and bright-field images of sorted sperm from caput (**c**) and corpus (**d**) epididymis of WT and dHET males. H3.3-mCherry and DNA staining confirm nuclear identity and purity of sorted sperm populations. Scale bars, 5 µm. **e,** Representative comet assay images illustrating intact, damaged, and fragmented DNA patterns in epididymal sperm. Arrowheads indicate DNA tails characteristic of damaged or fragmented nuclei. Scale bar, 100 µm. **f,** Quantification of comet assay patterns in caput and corpus sperm from WT and dHET males. Stacked bar plots show the relative frequencies of intact, damaged, and fragmented nuclei; numbers below bars indicate total cells scored.

**Supplementary Table 1.** Antibodies used in this study.

| Supplementary Table 1 \| Antibodies used in this study |  |  |  |  |
| --- | --- | --- | --- | --- |
| Antibodies | Company | Host species | Cat. No. | Application |
| anti-PRM1 | Briar Patch Biosciences | Mouse | Hup1N | IF and CUT&Tag |
| anti-PRM2 | Briar Patch Biosciences | Mouse | Hup2B | IF |
| Mouse IgG1 | Biolegend | Mouse | 400101 | CUT&Tag |
| anti-TNP1 | Proteintech | Rabbit | 17178-1-AP | IF |

**Supplementary Table 2.**
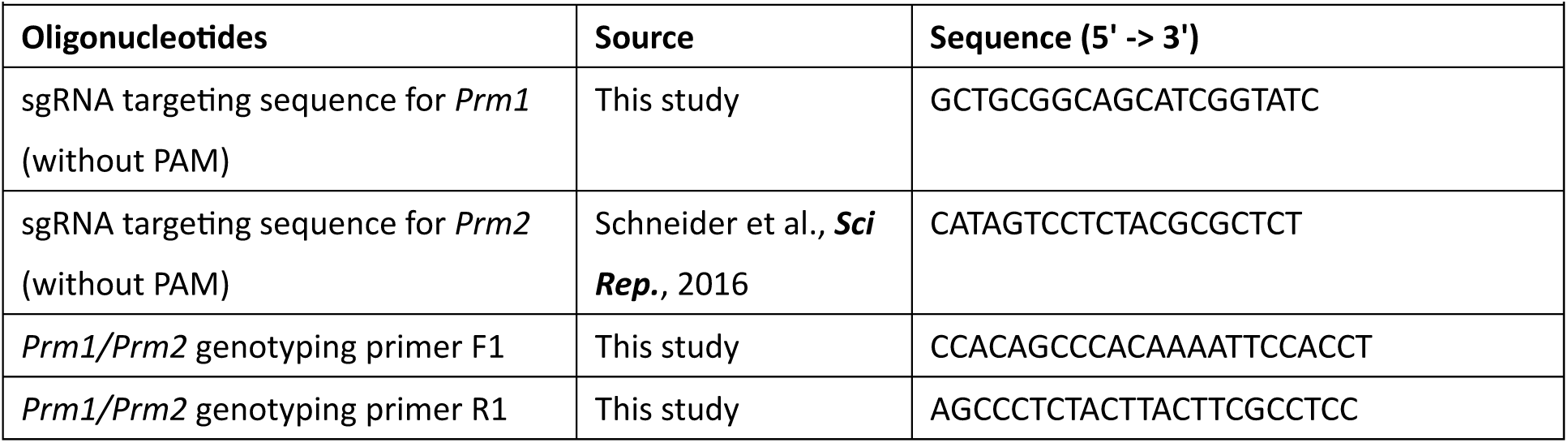
sgRNAs and primers.

## Notes

### Competing Interest Statement

The authors have declared no competing interest.

## References

1. Rathke, C., Baarends, W. M., Awe, S. & Renkawitz-Pohl, R. Chromatin dynamics during spermiogenesis. Biochim. Biophys. Acta 1839, 155–168 (2014).

2. Bao, J. & Bedford, M. T. Epigenetic regulation of the histone-to-protamine transition during spermiogenesis. Reproduction 151, R55–70 (2016).

3. Meistrich, M. L., Mohapatra, B., Shirley, C. R. & Zhao, M. Roles of transition nuclear proteins in spermiogenesis. Chromosoma 111, 483–488 (2003).

4. Arévalo, L., Esther Merges, G., Schneider, S. & Schorle, H. Protamines: lessons learned from mouse models. Reproduction 164, R57–R74 (2022).

5. Dhar, S., Thota, A. & Rao, M. R. S. Insights into role of bromodomain, testis-specific (Brdt) in acetylated histone H4-dependent chromatin remodeling in mammalian spermiogenesis. J. Biol. Chem. 287, 6387–6405 (2012).

6. Wang, X. et al. PHF7 is a novel histone H2A E3 ligase prior to histone-to-protamine exchange during spermiogenesis. Development 146, dev175547 (2019).

7. Kim, C. R. et al. PHF7 Modulates BRDT Stability and Histone-to-Protamine Exchange during Spermiogenesis. Cell Rep. 32, 107950 (2020).

8. Qian, M.-X. et al. Acetylation-mediated proteasomal degradation of core histones during DNA repair and spermatogenesis. Cell 153, 1012–1024 (2013).

9. Barral, S. et al. Histone variant H2A.L.2 guides transition protein-dependent protamine assembly in male germ cells. Mol. Cell 66, 89–101.e8 (2017).

10. Gill, M. E., Kohler, H. & Peters, A. H. F. M. Isolation of mouse germ cells by FACS using Hoechst 33342 and SYTO16 double staining. Methods Mol. Biol. 2770, 53–62 (2024).

11. Battulin, N. et al. Comparison of the three-dimensional organization of sperm and fibroblast genomes using the Hi-C approach. Genome Biol. 16, 77 (2015).

12. Jung, Y. H. et al. Chromatin States in Mouse Sperm Correlate with Embryonic and Adult Regulatory Landscapes. Cell Rep. 18, 1366–1382 (2017).

13. Chen, X. et al. Key role for CTCF in establishing chromatin structure in human embryos. Nature 576, 306–310 (2019).

14. Vara, C. et al. Three-Dimensional Genomic Structure and Cohesin Occupancy Correlate with Transcriptional Activity during Spermatogenesis. Cell Rep. 28, 352–367.e9 (2019).

15. Jessberger, G. et al. Cohesin and CTCF do not assemble TADs in Xenopus sperm and male pronuclei. Genome Res. 33, 2094–2107 (2023).

16. Yin, Q. et al. Revisiting chromatin packaging in mouse sperm. Genome Res. 33, 2079–2093 (2023).

17. Xu, H. et al. Three-dimensional genome structures of single mammalian sperm. Nat. Commun. 16, 3805 (2025).

18. Fujiwara, Y. et al. Isolation of stage-specific spermatogenic cells by dynamic histone incorporation and removal in spermatogenesis. Cytometry A 105, 297–307 (2024).

19. Alavattam, K. G. et al. Attenuated chromatin compartmentalization in meiosis and its maturation in sperm development. Nat. Struct. Mol. Biol. 26, 175–184 (2019).

20. Lin, C.-Y. et al. Human X-linked intellectual disability factor CUL4B is required for post-meiotic sperm development and male fertility. Sci. Rep. 6, 20227 (2016).

21. Schneider, S. et al. Protamine-2 deficiency initiates a reactive oxygen species (ROS)-mediated destruction cascade during epididymal sperm maturation in mice. Cells 9, 1789 (2020).

22. Merges, G. E. et al. Loss of Prm1 leads to defective chromatin protamination, impaired PRM2 processing, reduced sperm motility and subfertility in male mice. Development (2022) doi:10.1242/dev.200330.

23. Kaneko, S., Yoshida, J., Ishikawa, H. & Takamatsu, K. Single-cell pulsed-field gel electrophoresis to detect the early stage of DNA fragmentation in human sperm nuclei. PLoS One 7, e42257 (2012).

24. Hammoud, S. S. et al. Distinctive chromatin in human sperm packages genes for embryo development. Nature 460, 473–478 (2009).

25. Erkek, S. et al. Molecular determinants of nucleosome retention at CpG-rich sequences in mouse spermatozoa. Nat. Struct. Mol. Biol. 20, 868–875 (2013).

26. Hammoud, S. S. et al. Chromatin and transcription transitions of mammalian adult germline stem cells and spermatogenesis. Cell Stem Cell 15, 239–253 (2014).

27. Carone, B. R. et al. High-resolution mapping of chromatin packaging in mouse embryonic stem cells and sperm. Dev. Cell 30, 11–22 (2014).

28. Yamaguchi, K. et al. Re-evaluating the Localization of Sperm-Retained Histones Revealed the Modification-Dependent Accumulation in Specific Genome Regions. Cell Rep. 23, 3920–3932 (2018).

29. Luense, L. J. et al. Gcn5-Mediated Histone Acetylation Governs Nucleosome Dynamics in Spermiogenesis. Dev. Cell 51, 745–758.e6 (2019).

30. Gupta, S., Aggarwal, S. & Munde, M. New insights into the role of ligand-binding modes in GC-DNA condensation through thermodynamic and spectroscopic studies. ACS Omega 8, 4554–4565 (2023).

31. Guelen, L. et al. Domain organization of human chromosomes revealed by mapping of nuclear lamina interactions. Nature 453, 948–951 (2008).

32. van Steensel, B. & Belmont, A. S. Lamina-associated domains: Links with chromosome architecture, heterochromatin, and gene repression. Cell 169, 780–791 (2017).

33. Lieberman-Aiden, E. et al. Comprehensive mapping of long-range interactions reveals folding principles of the human genome. Science 326, 289–293 (2009).

34. Pereira, C. D., Serrano, J. B., Martins, F., da Cruz E Silva, O. A. B. & Rebelo, S. Nuclear envelope dynamics during mammalian spermatogenesis: new insights on male fertility. Biol. Rev. Camb. Philos. Soc. 94, 1195–1219 (2019).

35. Okada, Y. Sperm chromatin condensation: epigenetic mechanisms to compact the genome and spatiotemporal regulation from inside and outside the nucleus. Genes Genet. Syst. 97, 41–53 (2022).

36. Yu, Y. E. et al. Abnormal spermatogenesis and reduced fertility in transition nuclear protein 1-deficient mice. Proc. Natl. Acad. Sci. U. S. A. 97, 4683–4688 (2000).

37. Zhao, M. et al. Targeted disruption of the transition protein 2 gene affects sperm chromatin structure and reduces fertility in mice. Mol. Cell. Biol. 21, 7243–7255 (2001).

38. Shirley, C. R., Hayashi, S., Mounsey, S., Yanagimachi, R. & Meistrich, M. L. Abnormalities and reduced reproductive potential of sperm from Tnp1- and Tnp2-null double mutant mice. Biol. Reprod. 71, 1220–1229 (2004).

39. Zhao, M. et al. Transition nuclear proteins are required for normal chromatin condensation and functional sperm development. Genesis 38, 200–213 (2004).

40. Namekawa, S. H. et al. Postmeiotic sex chromatin in the male germline of mice. Curr. Biol. 16, 660–667 (2006).

41. Hada, M., Masuda, K., Yamaguchi, K., Shirahige, K. & Okada, Y. Identification of a variant-specific phosphorylation of TH2A during spermiogenesis. Sci. Rep. 7, 46228 (2017).

42. Tower, C. A., et al. Maternal CENP-C restores centromere symmetry in mammalian zygotes to ensure proper chromosome segregation. bioRxivorg (2025) doi:10.1101/2025.07.23.666394.

43. Makino, Y. et al. Generation of a dual-color reporter mouse line to monitor spermatogenesis in vivo. Front Cell Dev Biol 2, 30 (2014).

44. Corces, M. R. et al. An improved ATAC-seq protocol reduces background and enables interrogation of frozen tissues. Nat. Methods 14, 959–962 (2017).

45. Yamanaka, S., Kishi, Y. & Siomi, H. ATAC-seq method applied to embryonic germ cells and neural stem cells from mouse: Practical tips and modifications. in Epigenetics Methods (ed. Tollefsbol, T.) vol. 18 371–386 (Elsevier, 2020).

46. Fujiwara, Y. et al. Preparation of optimized concanavalin A-conjugated Dynabeads® magnetic beads for CUT&Tag. PLoS One 16, e0259846 (2021).

